# Selective mRNA translation determines adaptative mutability of melanoma cells to anti-BRAF/MEK combination therapy

**DOI:** 10.1101/2025.09.02.673634

**Authors:** Lucilla Fabbri, Lucie Lagadec, Eva Guérin, Hélène Lecourt, Dorothée Baille, Laetitia Besse, Cédric Messaoudi, Laurent Désaubry, Hussein Abou-Hamdan, Sévérine Roy, Bérangère Lombard, Damarys Loew, Jean-Yves Scoazec, Caroline Robert, Stéphan Vagner

**Affiliations:** Institut Curie, PSL Research University, CNRS UMR3348, INSERM U1278, Orsay, France; Université Paris-Saclay, CNRS UMR3348, INSERM U1278, Orsay, France; Equipe labellisée Ligue Nationale contre le Cancer, Orsay, France; INSERM U981, Gustave Roussy Cancer Campus, Villejuif, France; Université Paris-Sud, Université Paris-Saclay, Kremlin-Bicêtre, France; Dermato-Oncology, Gustave Roussy Cancer Campus, Villejuif, France; Institut Curie, PSL University, CNRS UAR2016, Inserm US43, Université Paris-Saclay, Multimodal Imaging Center, Orsay, France; Center of Research in Biomedicine of Strasbourg, Regenerative Nanomedicine (UMR 1260), INSERM, University of Strasbourg, Strasbourg, France; Institut Curie, PSL University, CurieCoreTech Mass Spectrometry Proteomics, Paris, France; Institut Gustave Roussy, INSERM U1015, Bâtiment de Médecine Moléculaire 114 rue Edouard Vaillant, 94800 Villejuif, France

## Abstract

During their inevitable evolution towards acquired resistance to anti-cancer targeted therapies, cancer cells adopt distinct gene expression profiles that allow them to transiently adapt to and tolerate the treatment. Similar to bacterial cells that transiently tolerate antibiotics, cancer cells surviving therapy can increase their mutation rate, enhancing the likelihood of acquiring resistance-conferring mutations and evolving into resistant cells. This adaptive mutability has been linked to transcriptional reprogramming of DNA damage repair mechanisms and effective therapeutic strategies to target such mechanisms are currently lacking. Here we show that translational control mediates the adaptive mutability of melanoma drug-tolerant cells by regulating the translation of the error-prone non-homologous end joining (NHEJ) component *53BP1*. The specific inhibition of 5’UTR-driven *53BP1* mRNA translation was sufficient to impair NHEJ and mutability. We found that the eIF4A RNA helicase, a key component of the eIF4F translation complex, regulates *53BP1* mRNA translation. Consequently, targeting the eIF4A with two small molecule inhibitors significantly delays the acquisition of resistance to combination of BRAF and MEK inhibitors (BRAFi/MEKi) in BRAF^V600^-mutant melanoma xenograft models and cell lines by reducing the mutability of drug-tolerant cells. Our results demonstrate that a standard-of-care therapy for melanoma, by engaging non-genetic adaptation driven at the translational level, contributes to the evolution of drug-tolerant melanoma cells toward acquired resistance.

## INTRODUCTION

Targeted therapy combining BRAF and MEK inhibitors (BRAFi/MEKi) has revolutionized the treatment of metastatic melanoma harboring the BRAF^V600^ mutation, which is present in approximately 50% of patients (1,2). Despite the initial clinical benefits, which achieve response rates of up to 70%, most patients eventually experience disease progression within several months due to the development of acquired resistance (3,4).

In addition to genetic mutations, recent findings have revealed the existence of non-genetic mechanisms that underpin drug tolerance in a subset of cancer cells and drive evolution towards resistance (5–7). Although the drug-tolerant state may initially be reversible, prolonged drug exposure can lead to increased mutation rates in drug-tolerant cells, ultimately promoting the emergence of clones characterized by stable, genetic resistance and cancer relapse (8–10). Such adaptive mutability has been shown to be threatened by accumulating DNA double-strand breaks (DSBs), even upon exposure to nongenotoxic therapies (9–12). In addition to the error-free homology-directed repair (HDR) pathway, which is restricted to the S and G2 phases of the cell cycle, DSB repair is also mediated by the non-homologous end joining (NHEJ) pathway (13). Unlike HDR, NHEJ operates throughout all phases of the cell cycle but is error-prone, potentially leading to deleterious mutations or deletions that may contribute to adaptive mutability. In melanoma, BRAFi/MEKi therapy has been proposed to induce transcriptional rewiring of DNA repair genes, leading to the suppression of error-free pathways, such HDR, and the emergence of a BRCAness-like state (14,15) Additionally, recent findings implicated the NHEJ pathway in the acquisition of genetic instability of melanoma following targeted therapy, promoting the evolution of acquired resistance (16).

Dynamic and transient changes in gene expression, through epigenetic, transcriptional or translational reprogramming, have been implicated in fostering cellular plasticity and enabling melanoma cells to adapt to therapy (17–20). Adaptive translational reprogramming has been previously implicated in mediating melanoma tolerance and acquisition of resistance to targeted therapies (20–25). We demonstrated that the emergence of non-genetic drug-tolerant melanoma cells is linked to eIF4F-mediated mRNA translation (20). This complex, consisting of the eIF4E cap-binding protein, the eIF4G scaffolding protein, and the eIF4A RNA helicase, binds to the cap structure at the 5’-end of mRNAs to mediate cap-dependent translation (26). Although the persistent formation of the eIF4F translation initiation complex has been associated with stable genetic resistance to BRAFi/MEKi therapy in BRAF^V600^-mutant melanoma (22), its potential role in driving the evolution of drug-tolerant cells toward acquired resistance through adaptive mutability remains unexplored.

Here we show that, in melanoma drug-tolerant cells, the adaptive mRNA translation reprogramming occurring following targeted therapy enhances the translation of *53BP1* mRNA, which encodes a crucial NHEJ promoter (27). This correlates with enhanced 53BP1-dependent mutagenesis and a shift toward NHEJ-directed repair. Notably, CRISPR/Cas9-mediated partial deletion of the *53BP1* 5′ UTR in drug-tolerant cells reduces NHEJ activity and impairs mutability, underscoring the importance of translational control.

We identify eIF4A as a key regulator of *53BP1* mRNA translation. Treatment with the clinically advanced eIF4A inhibitor eFT226 significantly impairs *53BP1* mRNA translation, NHEJ activity, and the mutability of drug-tolerant cells. Furthermore, co-treatment with BRAFi/MEKi and either eFT226 or a newly developed, selective eIF4A inhibitor, RBX0901, delays the onset of acquired resistance in mouse xenograft models. Collectively, our findings reveal that translational regulation of *53BP1* mRNA is associated with the transition from a reversible drug-tolerant state to permanent genetic resistance by promoting NHEJ and mutability. Targeting this regulatory axis by inhibiting eIF4A may offer a potential therapeutic strategy to limit tumor evolution and recurrence.

## RESULTS

### Targeted therapy increases the expression of the NHEJ protein 53BP1

Quantitative proteomic analysis performed on BRAF^V600E^-mutated A375 melanoma cells that survived 3 days-exposure to high-dose BRAF/MEK inhibitors (BRAFi/MEKi) revealed a profound reprogramming of the proteome compared to untreated cells (DMSO), with DNA repair–related pathways ranking among the top 10 most dysregulated ones (Fig 1A,B). HDR pathway was markedly downregulated in BRAFi/MEKi-surviving A375 cells, supporting the hypothesis that targeted therapy promotes a BRCAness-like phenotype (Fig. S1A). By employing another BRAF ^V600E^ mutated cell line sensitive to MAPK pathway inhibition (28), we observed an increase in γH2AX levels in A375 and M249 cells that survived BRAFi/MEKi, (Fig. 1C,D) suggesting potential DNA damage induced by the treatment. Such increase was specifically associated with drug exposure, as re-culturing therapy-surviving cells in drug-free media for 9 days decreased the quantity of the protein to levels observed in untreated cells (Fig 1C,D), thereby suggesting that therapy-induced DNA damage is reversible upon drug withdrawal.

**Figure 1:**
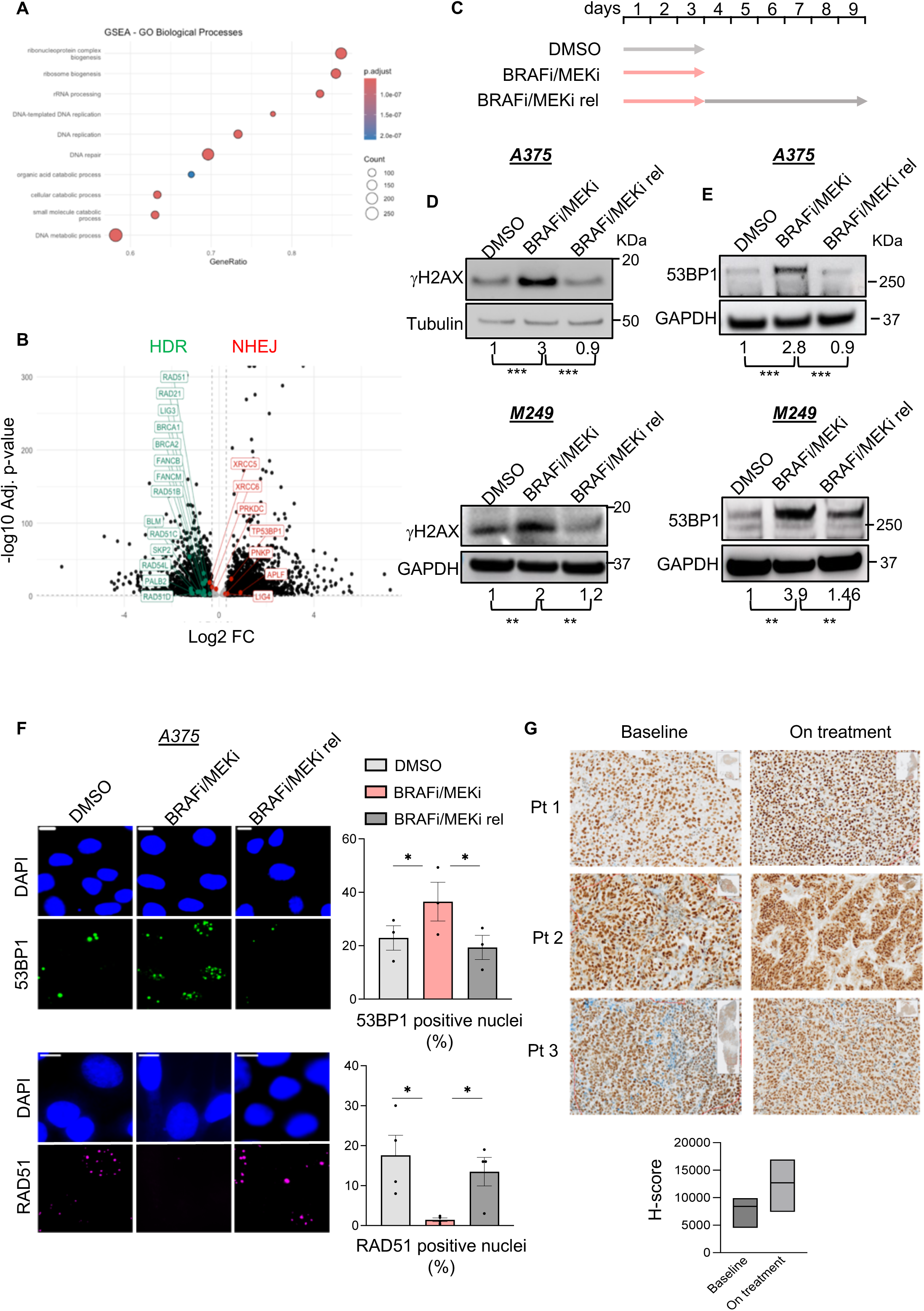
Targeted therapy increases the expression of the NHEJ protein 53BP1. **A**, Dot plot showing significantly enriched gene sets of differentially expressed proteins in BRAF^V600E^-mutated A375 melanoma cells surviving BRAFi/MEKi therapy (BRAFi/MEKi) versus A375 cells treated with DMSO using GSEA. The color of the bubbles represents the p-value adjusted, and the size of the bubbles represents the number of enriched genes from each pathway. **B**, Volcano plots showing *P* values adjusted (−log_10_) of versus the log_2_ fold change (FC) of the proteomics analysis of BRAFi/MEKi-treated A375 cells versus DMSO-treated cells (*n* = 3); non-significant proteins are shown in gray and proteins involved in HDR and NHEJ pathway are highlighted in green and red, respectively and key proteins are labeled (*n* = 3). **C,** Schematic representation of cell treatments used for subsequent analysis. Cells were treated with DMSO (control) or with targeted therapy (BRAFi/MEKi) for 3 days. Cells that survived therapy were re-cultured in drug-free media for an additional 9 days (BRAFi/MEKi rel). **D**, Western blot illustrating γH2AX levels in A375 and M249 cells treated with DMSO, in cells surviving targeted therapy (BRAFi/MEKi) or in cells released from the drugs for 9 days (BRAFi/MEKi rel). GAPDH or Tubulin serve as a loading control, and the relative quantification from 3 independent experiments is indicated (** p ≤ 0.01). **E**, Western blot illustrating 53BP1 levels in A375 and M249 cells treated with DMSO, in cells surviving targeted therapy (BRAFi/MEKi) or in cells released from the drugs for 9 days (BRAFi/MEKi rel). GAPDH serves as a loading control, and the relative quantification from 3 independent experiments is indicated (** p ≤ 0.01, *** p ≤ 0.001). **F**, (top) Representative images of immunofluorescence in A375 control (DMSO), drug tolerant cells (BRAFi/MEKi) and drug tolerant cells cells re-cultured released from targeted therapy for 9 days (BRAFi/MEKi rel) stained for 53BP1 (green) and nuclei (DAPI, blue). (bottom) Representative images of immunofluorescence in cells stained for RAD51 (magenta) and nuclei (DAPI, blue). Scale bar, 10 μm. The quantification of the signal analyzed by percentage of cells with ≥ 10 foci/nucleus is reported and the data shown represent the mean ± SEM from 3 independent experiments. p-values were calculated by paired, two-tailed Student’s t-test (* p ≤ 0.05). **G**, (top) Representative images of immunohistochemistry for 53BP1 in melanoma patient biopsies collected at baseline or ON treatment. Analysis of the average score of 53BP1 staining between the two conditions is shown (bottom).

It is known that NHEJ can compensate for defects in HDR (14,29). Our proteomic analysis showed that protein levels of NHEJ-related factors were either maintained or increased following BRAFi/MEKi exposure (Fig. 1B and Fig. S1B). To identify potential mediators of NHEJ with increased mRNA translation following BRAFi/MEKi-targeted therapy, we reanalyzed our previous genome-wide polysome profiling data (20). Among the translationally upregulated mRNAs, we selected the one encoding the key NHEJ mediator 53BP1. 53BP1 is a chromatin binding protein that regulate DSB repair pathway choice by promoting canonical NHEJ-mediated DSB repair (27). BRAFi/MEKi treatment led to a reversible increase in 53BP1 protein levels, as analyzed by western blot (Fig. 1E), without changes in *53BP1* mRNA levels (Fig. S1C).

Analysis of the nuclear recruitment of 53BP1 and RAD51, a key effector of HDR, revealed that BRAFi/MEKi treatment increased the number of nuclei positive for 53BP1, while decreasing the ones positive for RAD51 in both A375 and M249 cells (> 10 foci/nucleus) (Fig. 1F and **Fig. S1D**). Upon drug withdrawal, nuclear recruitment patterns of these proteins returned to levels similar to those observed in untreated cells (Fig. 1F and Fig. S1D) indicating that, in the two tested BRAF^V600E^ mutated melanoma cell lines, targeted therapy reprograms DNA repair by reversibly shifting the balance from HDR to NHEJ.

We next investigated 53BP1 protein levels in a pilot analysis of matched clinical biopsy specimens from 3 melanoma patients collected at baseline and upon BRAFi/MEKi exposure. Samples collected on treatment showed a trend towards increased 53BP1 protein levels in compared to baseline (Fig. 1G). While this observation is based on a limited sample set, it suggests that the induction of 53BP1 upon targeted therapy could extend as well to clinical specimen. Altogether, these results show that 53BP1 expression is modulated upon targeted therapy.

### 5’UTR-dependent *53BP1* mRNA translation is induced upon targeted therapy

To next evaluate the effect of BRAFi/MEKi-targeted therapy on *53BP1* mRNA translation, we isolated actively translating ribosomes following short L-azidohomoalanine (AHA) pulse and click chemistry, and their associated transcripts (Fig. 2A). A375 and M249 cells that survived 3 days of BRAFi/MEKi-targeted therapy exhibited increased association of *53BP1* mRNA with active ribosomes compared to untreated cells (Fig. 2B). The increase in *53BP1* mRNA translation mirrored the kinetic of protein expression observed in Fig. 1E, returning to levels observed in untreated cells after 9 days of drug withdrawal (Fig. 2B). Given the observed increase in 53BP1 protein levels in a subset of drug-tolerant cells in Fig. 1F, we intended to measure protein synthesis at a single cell level by using the recently described ribosome-bound mRNA mapping (RIBOmap) assay (30). This approach detects ribosome-bound mRNAs through a proximity-based probe system (Fig. 2C). When in proximity, the three probes generate a DNA amplicon through rolling-cicle amplification, indicative of active translation. To detect actively translated mRNAs, we used a fluorescent probe complementary to the DNA amplicon. Translating *53BP1* mRNAs decreased upon *53BP1* downregulation (Fig. 2D,E, Fig. S2A,B), validating the specificity of the assay. We found that the increase of *53BP1* mRNAs association with active ribosomes occurred preferentially in a subset of drug-tolerant cells (Fig. 2F,G, Fig. S2C). Consistently, a subpopulation of drug-tolerant cells exhibited increased translation of IQGAP1 mRNA, which we previously demonstrated to be another translationally regulated mRNA in drug-tolerant cells (Fig. S2D) (20). Combined analysis of translating *53BP1* and *IQGAP1* mRNAs in A375 drug tolerant cells using the RiboMap assay (Fig. 2H) revealed a significant correlation between the translation of *53BP1* and *IQGAP1* mRNAs (Fig. 2I and Fig. S2E), with a Pearson correlation coefficient ranging from 0.62 to 0.8, suggesting a mechanism of translational co-regulation. Consistently, a single cell multiplexed imaging approach by co-detection by indexing (CODEX) indicated that a subpopulation of drug-tolerant cells exhibited the highest staining intensity for 53BP1, IQGAP1 and all other 4 selected markers (CCSER2, HIPK2, ANK2 and APC), whose expression was previously characterized as translationally regulated in drug-tolerant cells (20). These results suggest that BRAFi/MEKi-induced selective translational regulation promotes their coordinated expression in this subset of cells. In contrast, such co-expression was not observed for the other markers whose expression was shown to be inducedat the transcriptional level (e.g. SLIT2, CD36) (18) (Fig. 2J,K, Fig. S3).

**Figure 2:**
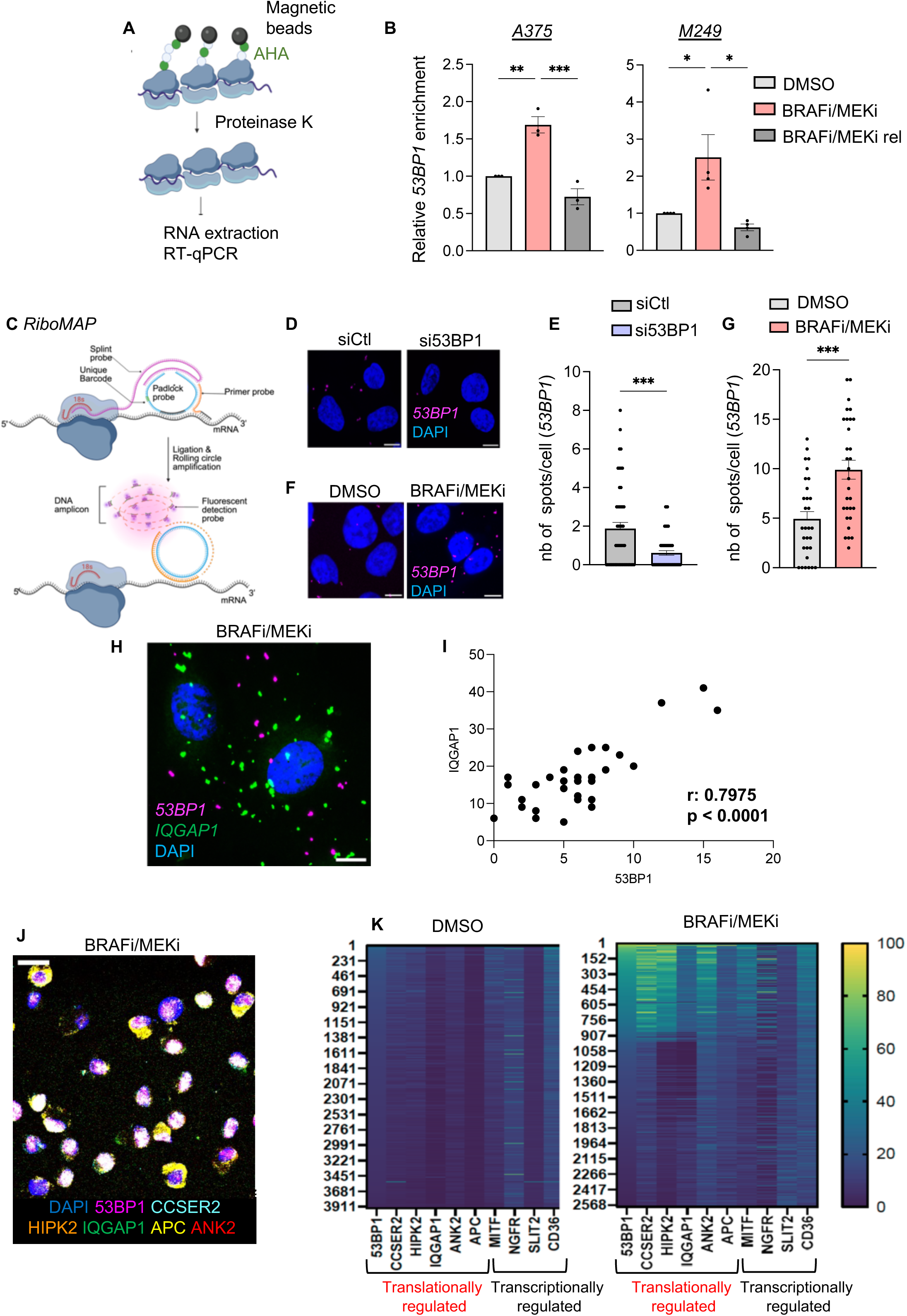
5’UTR-dependent *53BP1* mRNA translation is induced upon targeted therapy. **A**, Schematic representation of the protocol used for the isolation of actively translating ribosomes and their associated transcripts. After a short L-azidohomoalanine (AHA) pulse, newly synthesized AHA-labelled peptides are used to isolate active ribosome complexes through chemical interactions with magnetic beads. Bead-bound complexes are then treated with Proteinase K and the purified RNA is used for RT-qPCR. **B**, Quantification of *53BP1* mRNA enrichment in active ribosomes performed using RT-qPCR in A375 and M249 control cells (DMSO), in cells surviving targeted therapy (BRAFi/MEKi) and in cells released from the drugs for 9 days (BRAFi/MEKi rel). Data represent the mean ± SEM from 3 (A375) or 4 (M249) independent experiments (* p ≤ 0.05, ** p ≤ 0.01, *** p ≤ 0.001). **C**, Schematic of the ribosome-bound mRNA mapping (RIBOmap) used to detect translating mRNAs. RIBOmap relies on the use of a tri-probe set: (1) a primer probe that hybridizes to the target mRNAs, (2) a splint DNA probe that hybridizes with the ribosomal 18S RNA, and (3) a padlock probe. When in proximity, the tri-probes produce DNA amplification signals corresponding to active mRNA translation. **D**, Representative confocal images of translating 53*BP1* mRNAs (magenta) in A375 cells transfected with control siRNA (siCtl) or with siRNA targeting *53BP1* (si53BP1) for 48 hours. Nuclei are stained with DAPI (blue). Scale bar: 10 µm. **E**, Quantification of translating *53BP1* mRNAs (spots) in the condition described in D. Data shown represent the mean ± SEM of the number of spots/cells quantified in 1 independent experiment (see also Fig. S2B for the other biological replicates) (*** p ≤ 0.001). **F**, Representative confocal images of translating 53*BP1* mRNAs (magenta) assessed by RiboMap assay in A375 control cells (DMSO) or in cells surviving targeted therapy (BRAFi/MEKi). Nuclei are stained with DAPI (blue). Scale bar: 10 µm. **G**, Quantification of translating *53BP1* mRNAs (spots) in the condition described in F. Data shown represent the mean ± SEM of the number of spots/cells quantified in 1 independent experiment(see also Fig. S2C for the other biological replicates) (*** p ≤ 0.001). **H**, Representative confocal images of translating *53BP1* (magenta) and IQGAP*1* mRNAs (green) assessed by RiboMap assay in A375 cells surviving targeted therapy. Nuclei are stained with DAPI (blue). Scale bar: 10 µm. **I**, Scatterplot illustrating the correlation between translating *53BP1* mRNAs and *IQGAP1* mRNAs in A375 drug tolerant cells. Pearson correlation coefficient (r) and p-value are reported. The data shown are the results of 1 biological replicate (see Supplementary Figure S2E for the results obtained in other biological replicates). **J**, Multiplex immunostaining (CODEX) showing the co-expression of 53BP1, CCSER2, HIPK2, IQGAP1, APC, and ANK2 in A375 cells. Scale bar 20 µm. **K**, Heatmap generated from multiplexed imaging analysis, displaying single-cell mean intensity values (rows) for protein expression of selected differentially expressed markers (columns), regulated at either the translational or transcriptional level in control cells (DMSO) and drug-tolerant cells (BRAFi/MEKi).

Given the essential role of the 5’UTR in regulating translation of mRNAs (31), we next investigated the role of *53BP1* 5’UTR in regulating its translation. To this purpose, we used a luciferase expression reporter construct in which the *53BP1* 5’UTR was cloned upstream of the Renilla (LucR) open reading frame (Fig 3A). Firefly expression - LucF - from the same plasmid served as a control for transfection/expression. We monitored the luciferase activities in transfected A375 and M249 cells upon BRAFi/MEKi treatment. We observed that BRAFi/MEKi induced an increase in the LucR/LucF activity ratio in cells transfected with the *53BP1* 5′ UTR-containing reporter upon BRAFi/MEKi treatment, which returned to the levels observed in untreated cells after 9 days of drug withdrawal (Fig. 3B). This increase in LucR activity upon BRAFi/MEKi reflected enhanced translation, as the LucR mRNA levels remained unchanged relative to the LucF mRNA (Fig. 3C). These data indicate that *53BP1* 5’UTR is sufficient to mediate the increase in *53BP1* mRNA translation upon BRAFi/MEKi treatment.

**Figure 3:**
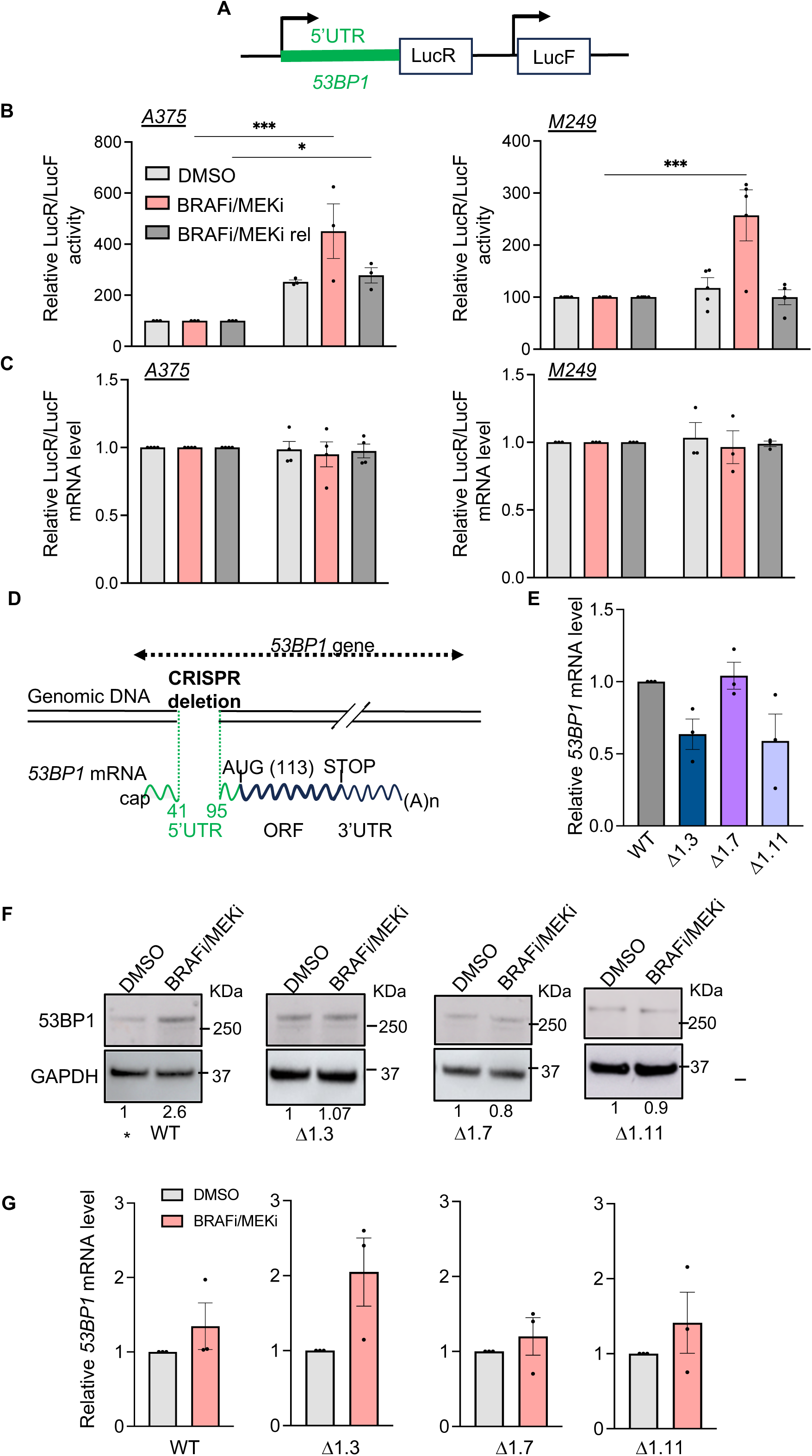
5’UTR-dependent *53BP1* mRNA translation is induced upon targeted therapy. **A**, Schematic of the 53BP1 5′ UTR-containing luciferase reporter. **B**, Quantification of the luciferase assay performed in A375 and M249 cells treated with DMSO or BRAFi/MEKi for 3 days, and in cells released from BRAFi/MEKi treatment (BRAFi/MEKi rel). Renilla (LucR) activity was measured 48h after transfection and the activity of the Firefly luciferase (LucF) was used as control of transfection. The data shown are normalized to the Empty vector (without *53BP1* 5’UTR) and represent the mean ± SEM from 3 (A375) or 4 (M249) independent experiments (* p ≤ 0.05, ** p ≤ 0.01, *** p ≤ 0.001). **C**, RT-qPCR quantification of the LucR/LucF mRNA level from the experiment in B. The data shown are normalized to the Empty vector. **D**, Schematic representation of genomic deletion by CRISPR/Cas9 genome editing leading to deleted 53BP1 5’UTR (from nucleotide 41 to 95) in A375 cells. **E**, RT-qPCR quantification of *53BP1* mRNA level in A375 WT cells or in A375 Δ 5’UTR clones. Data represent the mean ± SEM from 3 independent experiments. **F**, Western blot illustrating 53BP1 protein levels in A375 WT cells and A375 Δ5’UTR clones. For each cell line, control cells (DMSO) or drug tolerant cells that survived 3 days of targeted therapy (BRAFi/MEKi) were analyzed for 53BP1 protein levels. GAPDH serves as a loading control and the relative quantification from 4 independent experiments is indicated (** p ≤ 0.01). **G**, RT-qPCR quantification of *53BP1* mRNA level in A375 WT cells or in A375 Δ 5’UTR clones. For each cell line, control cells (DMSO) or drug tolerant cells that survived 3 days of targeted therapy (BRAFi/MEKi) were analyzed for *53BP1* mRNA level. Data represent the mean ± SEM from 3 independent experiments.

We next endogenously deleted a genomic region within the first exon of *53BP1* corresponding to nucleotides 41–95 of the *53BP1* 5′ UTR in A375 cells (Δ5’UTR) (Fig. 3D). The 5’UTR deletion prevented the increase in *53BP1* mRNA translation in response to BRAFi/MEKi treatment, without affecting its translation in untreated cells (Fig. S4A). Analysis of *53BP1* transcript levels in 3 different clones indicated that the 5’UTR deletion did not affect promoter functions (Fig. 3E) but abrogated the increase in 53BP1 protein levels observed upon BRAFi/MEKi (Fig. 3F) without changing *53BP1* mRNA levels (Fig. 3G). Altogether, these data show that the *53BP1* 5′ UTR is necessary to mediate the translational upregulation in drug-tolerant cells.

### *53BP1* mRNA translation promotes NHEJ upon targeted therapy

Since 53BP1 is a key promoter of NHEJ, we next investigated whether the increase in NHEJ efficiency observed in drug-tolerant cells was a direct consequence of enhanced translational upregulation of *53BP1* mRNA. We first evaluated NHEJ activity of A375 cells by measuring random DNA integration into genomic DNA (32,33). This assay measures the NHEJ-dependent genomic integration of a GFP-expressing linearized plasmid containing a Neomycin resistance cassette and integration events are scored as number of colonies resistant to G418 (33). Supporting this shift, A375 cells tolerant to BRAFi/MEKi treatment exhibited increased NHEJ efficiency compared to untreated controls an effect that was abolished by siRNA-mediated depletion of 53BP1 (Fig. 4A). To assess whether the NHEJ bias induced by targeted therapy was associated with increased *53BP1* mRNA translation, we evaluate NHEJ efficiency in three 5’UTR-deleted clones, where the targeted therapy-induced translation regulation of *53BP1* mRNA was compromised as shown in Fig. 3.

**Figure 4:**
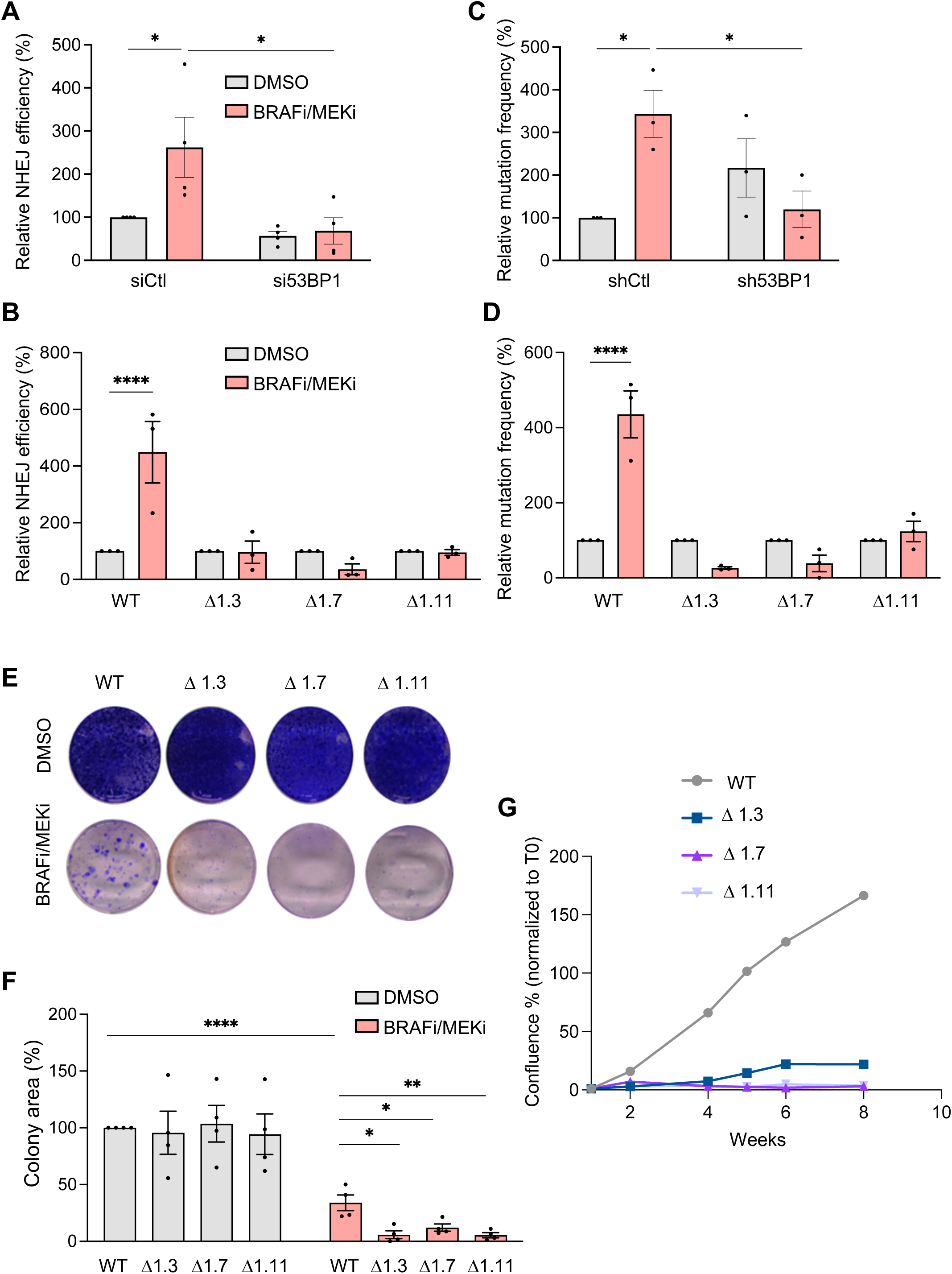
*53BP1* mRNA translation promotes NHEJ and mutability upon targeted therapy. **A**, Quantification of plasmid integration efficiencies of A375 control (DMSO) and BRAFi/MEKi drug tolerant cells (BRAFi/MEKi) transfected with control siRNA (siCtl) or siRNA targeting *53BP1* (si53BP1). The mean ± SEM from 4 independent experiments is shown (* p ≤ 0.05). Data were normalized to A375 control cells transfected with siCtl, which were set to 100%. **B**, Quantification of plasmid integration efficiencies of A375 WT cells and of 3 different A375 Δ 5’UTR clones following DMSO treatment or BRAFi/MEKi targeted therapy. The mean ± SEM from 3 independent experiments is shown (**** p ≤ 0.0001). Data were normalized to cells treated with DMSO, which were set to 100%. **C**, Relative HPRT mutation frequency for A375 cells stably expressing shCtl or sh53BP1, analysed in cells treated with DMSO or in BRAFi/MEKi drug tolerant cells (BRAFi/MEKi). Data represent the mean ± SEM from 3 independent experiments (* p ≤ 0.05) and are expressed as a fraction of the mutation frequency (%) observed in A375 cells expressing shCtl treated with DMSO. **D,** Relative HPRT mutation frequency analyzed in control (DMSO) or drug tolerant (BRAFi/MEKi) A375 WT cells or in 3 different A375 Δ 5’UTR clones. Data represent the mean ± SEM from 3 independent experiments (**** p ≤ 0.0001) and are expressed as a fraction of the mutation frequency (%) observed in control cells treated with DMSO. **E**, Colony assay performed 3 weeks after continuous treatment of A375 WT or Δ5’UTR drug tolerant cells (10000 cells) with DMSO or BRAFi/MEKi (100 nM/ 10 nM). **F**, Quantifications of the colony assay (mean ± SEM) from 4 independent experiments. p-values were calculated by unpaired, two-tailed Student’s t-test (*p ≤0.05, ** p ≤ 0.01, **** p ≤ 0.0001). **G**, Cell proliferation assay performed in A375 WT and Δ5’UTR drug tolerant cells. Cells surviving BRAFi/MEKi treatment were plated, and cell proliferation was monitored under continuous BRAFi/MEKi (100 nM/ 10 nM) treatment for 4 weeks (*n* = 1 biological experiment).

Similar to the effects of siRNA-mediated depletion of 53BP1, the enhanced NHEJ efficiency observed in A375 cells upon targeted therapy was abrogated in the 3 tested 5’UTR-deleted clones surviving BRAFi/MEKi-targeted therapy (Fig. 4B). Accordingly, these clones showed no targeted therapy-induced increase in 53BP1 positive cells (Fig. S4B,C). Consistent results were obtained across multiple clones, limiting potential off-target effects from CRISPR/Cas9-mediated deletion. These results indicate that the translational upregulation of *53BP1* induced by targeted therapy promotes NHEJ.

### *53BP1* mRNA translation promotes targeted therapy-induced mutability and the evolution towards acquired resistance

Recent studies showed that NHEJ-mediated repair plays a critical role in promoting genomic instability and the development of melanoma acquired resistance to targeted therapy (16). NHEJ-induced genomic instability may therefore increase the mutation rate in drug-tolerant cells, enhancing the likelihood of their evolution into genetically resistant clones (9,10,12). We thus investigated whether the 53BP1-mediated increase in NHEJ contributes to the induction of adaptive mutability in drug-tolerant melanoma cells. To this purpose, we analyzed the occurrence of genomic mutations upon BRAFi/MEKi by measuring loss-of-function mutations at the hypoxanthine-guanine phosphoribosyltransferase (HPRT) gene and consequent cells resistance to 6-thioguanine (34,35). In line with a role of 53BP1, shRNA-mediated knockdown of 53BP1, significantly reduced the enhanced mutation frequency observed in non-targeting shRNA control cells following BRAFi/MEKi-targeted therapy (Fig. 4C and Fig. S4D). To directly assess whether this increased mutability was driven by elevated *53BP1* mRNA translation, we evaluated mutation frequency at the *HPRT* locus in 3 different Δ5’UTR clones. In these clones, targeted therapy did not lead to an increase in mutation frequency compared to untreated cells (Fig. 4D). We next assessed whether the reduced mutability of drug-tolerant cells observed in Δ5’UTR clones was responsible for limiting the clonal capacity of drug-tolerant cells upon BRAFi/MEKi treatment. Compared to WT cells, drug-tolerant cells derived from Δ5′UTR clones exhibited a significant reduction in clonal emergence following continuous BRAFi/MEKi exposure for 3 weeks (Fig. 4E,F). By contrast, no differences in clonogenic capacity were observed between WT and Δ5′UTR cells under DMSO-treated control conditions. In line with this, monitoring of drug-tolerant cells proliferation over a two-month period revealed that drug-tolerant cells from Δ5′UTR clones displayed markedly impaired proliferative capacity compared to WT cells (Fig. 4G).

Together, these findings demonstrate that impairing 53BP1 translational regulation in drug-tolerant cells limits NHEJ capacity and the emergence of drug-tolerant-derived resistant and proliferative clones, thereby potentially overcoming acquired resistance.

### The use of the eIF4A inhibitor eFT226 prevents *53BP1* mRNA translation and adaptive mutability of drug-tolerant cells

Since we established a role for the *53BP1* 5′UTR in promoting increased mRNA translation in drug-tolerant cells, and given the previously demonstrated translational reprogramming in drug-tolerant cells mediated by the helicase eIF4A of eIF4F complex (20), we sought to assess the clinical relevance of this translational regulation by targeting the eIF4A. To this end, we employed the clinically advanced, flavagline-derivate, small-molecule inhibitor eFT226 (38). Similar to other flavaglines (39,40), eFT226 selectively represses the translation of mRNAs containing polypurine motifs in their 5′UTRs (38).

eFT226 specifically inhibited the increase in 53BP1 protein level observed in A375 drug-tolerant cells (Fig. 5A,B). This reduced 53BP1 protein expression in drug-tolerant cells corresponded to a reduced association of *53BP1* mRNA with active ribosomes following eFT226 treatment (Fig. 5C,D) indicating decreased translation. Consistently, in A375 drug-tolerant cells, eIF4A inhibition by eFT226 specifically decreased the translation of the 53BP1 5′ UTR-containing reporter but had no effect on the translation of a reporter mRNA containing no 5’UTR (“Empty”) (Fig. 5E) or the tubulin 5’UTR (Fig. S8), indicating that the *53BP1* 5’UTR is sufficient to observe both the BRAFi/MEKi or the eFT226 effects on translation.

**Figure 5:**
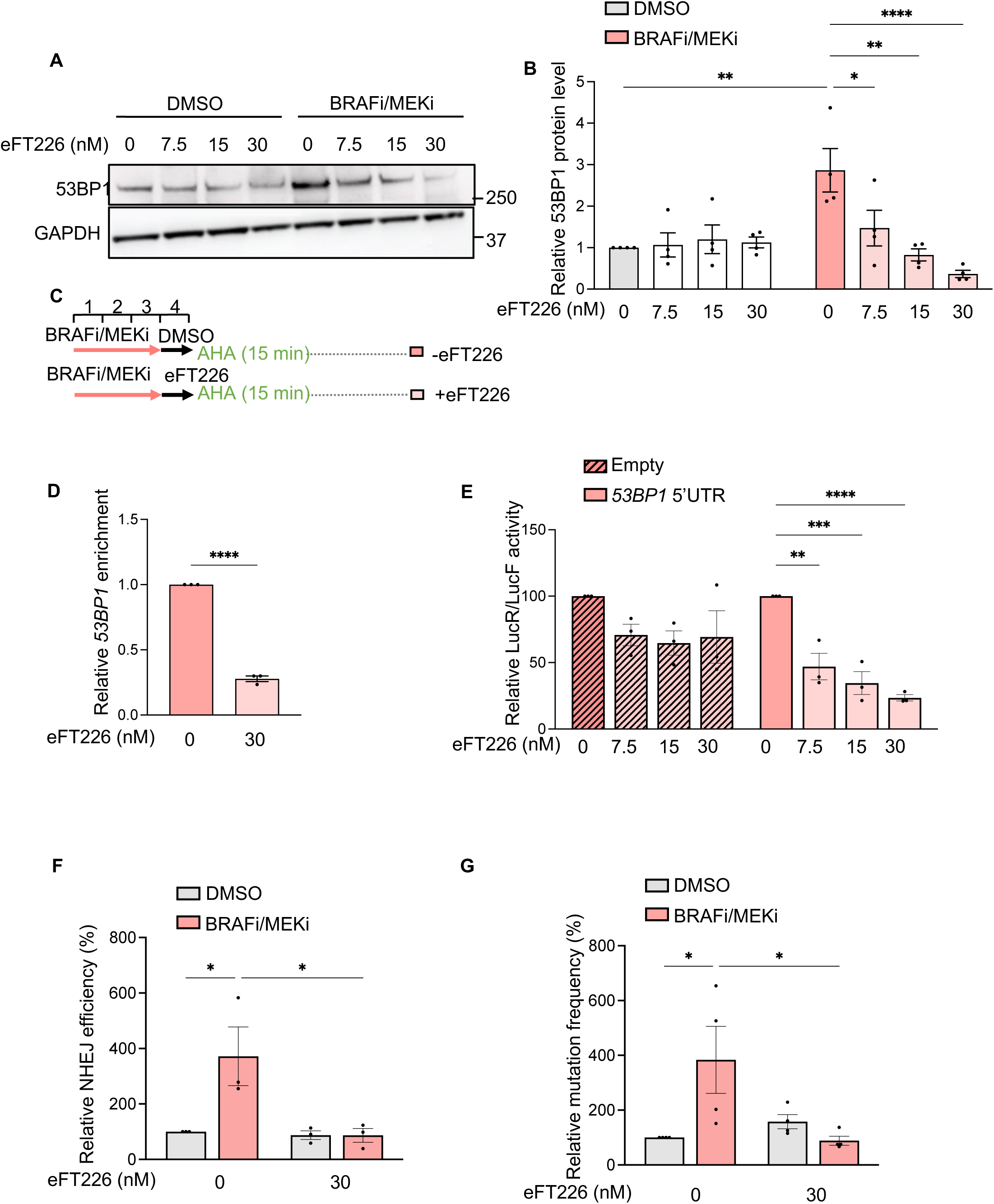
Targeting eIF4A with the use of eFT226 Inhibits *53BP1* mRNA translation and decreases NHEJ and mutability upon targeted therapy. **A**, Western blot illustrating 53BP1 protein levels in A375 control (DMSO) or drug tolerant cells (BRAFi/MEKi), treated with the indicated concentration of eFT226 for 24 hours. GAPDH serves as a loading control. **B**, Relative quantification of 53BP1 protein level in A375 cells treated as in A. Data represent the mean ± SEM from 4 independent experiments (* p ≤ 0.05, ** p ≤ 0.05, **** p ≤ 0.0001). **C**, Schematic representation of the protocol used for the isolation of actively translating ribosomes and their associated transcripts in A375 drug tolerant cells (surviving 3 days BRAFi/MEKi therapy) treated with DMSO or 30 nM of eFT226 for 24 hours. **D**, Quantification of *53BP1* mRNA enrichment in active ribosomes performed using RT-qPCR. Data represent the mean ± SEM from 3 independent experiments (**** p ≤ 0.0001). **E,** Quantification of the luciferase assay performed in A375 drug tolerant cells treated with the indicated concentrations of eFT226 for 12 hours. Renilla (LucR) activity was measured at the end of the eFT226 treatment, and the activity of the Firefly luciferase (LucF) was used as control of transfection. Data represent the effect of eFT226 on the luciferase activity of the reporter containing 53BP1 5’UTR or the Empty vector (without 53BP1 5’UTR) and are normalized to the eFT226 untreated conditions. The mean ± SEM from 3 independent experiments is reported (** p ≤ 0.01, *** p ≤ 0.001 **** p ≤ 0.0001). **F,** Quantification of plasmid integration efficiencies of A375 control (DMSO) and BRAFi/MEKi drug tolerant cells treated with or without eFT226 (30 nM for 24h). The mean ± SEM from 3 independent experiments is shown (* p ≤ 0.05). Data were normalized to A375 control cells treated with DMSO, which were set to 100%. **G,** Relative *HPRT* mutation frequency for A375 control cells treated with DMSO or BRAFi/MEKi drug tolerant cells, treated with or without eFT226 (30 nM for 24h). Data represent the mean ± SEM from 4 independent experiments (* p ≤ 0.05) and are expressed as a fraction of the mutation frequency (%) observed in A375 control cells treated with DMSO.

We next evaluated the NHEJ efficiency in A375 cells treated with eFT226 by measuring random DNA integration into genomic DNA (32,33). eFT226 reduced the enhanced NHEJ efficiency observed in cells surviving BRAFi/MEKi targeted therapy (Fig. 5F), which was associated with decreased 53BP1 nuclear recruitment (Fig. S5A,B). The reduced NHEJ efficiency of drug-tolerant cells by eIF4A inhibiton was accompanied by a decrease in the mutation frequency observed at the *HPRT* locus (Fig. 5G). These findings suggest that eIF4A inhibition reduces the BRAFi/MEKi targeted therapy-induced adaptive mutability in A375 cells by impairing NHEJ, thereby potentially delaying the development of stable resistance to BRAFi/MEKi targeted therapy.

### Combining BRAFi/MEKi targeted therapy with small molecule eIF4A inhibitors delays melanoma recurrence

To explore the therapeutic potential of targeting eIF4A-mediated *53BP1* mRNA translation in cancer cells transitioning from reversible non-genetic to irreversible genetic resistance to BRAFi/MEKi, we evaluated the efficacy of combining eFT226 with BRAFi/MEKi targeted therapy. We used an A375 melanoma xenograft model in which tumours initially respond to BRAFi/MEKi therapy but eventually relapse (41). Nude-athymic mice with established A375 orthotopic tumours (>150 mm^3^) were fed with a control chow or a chow supplemented with PLX4720 (BRAFi) and PD0325901 (MEKi) and underwent intraperitoneal injection with or without 1mg/kg of eFT226 twice a week. Notably, combinations were generally well tolerated for the duration of the study (Fig. S6). While BRAFi/MEKi triggered potent tumour regression with a complete disappearance of the tumor after 10-11 days of treament, eFT226 was ineffective (Fig. 6A). The combination of eFT226 with BRAFi/MEKi targeted therapy significantly delayed the development of resistance (Fig. 6A), with 10 out of 12 mice showing no relapse (tumor volume < 50 mm³) at the end of the experiment (Fig. 6B). This was associated with a prolonged median progression-free survival (PFS) compared to BRAFi/MEKi treatment alone (Fig. 6C).

**Figure 6:**
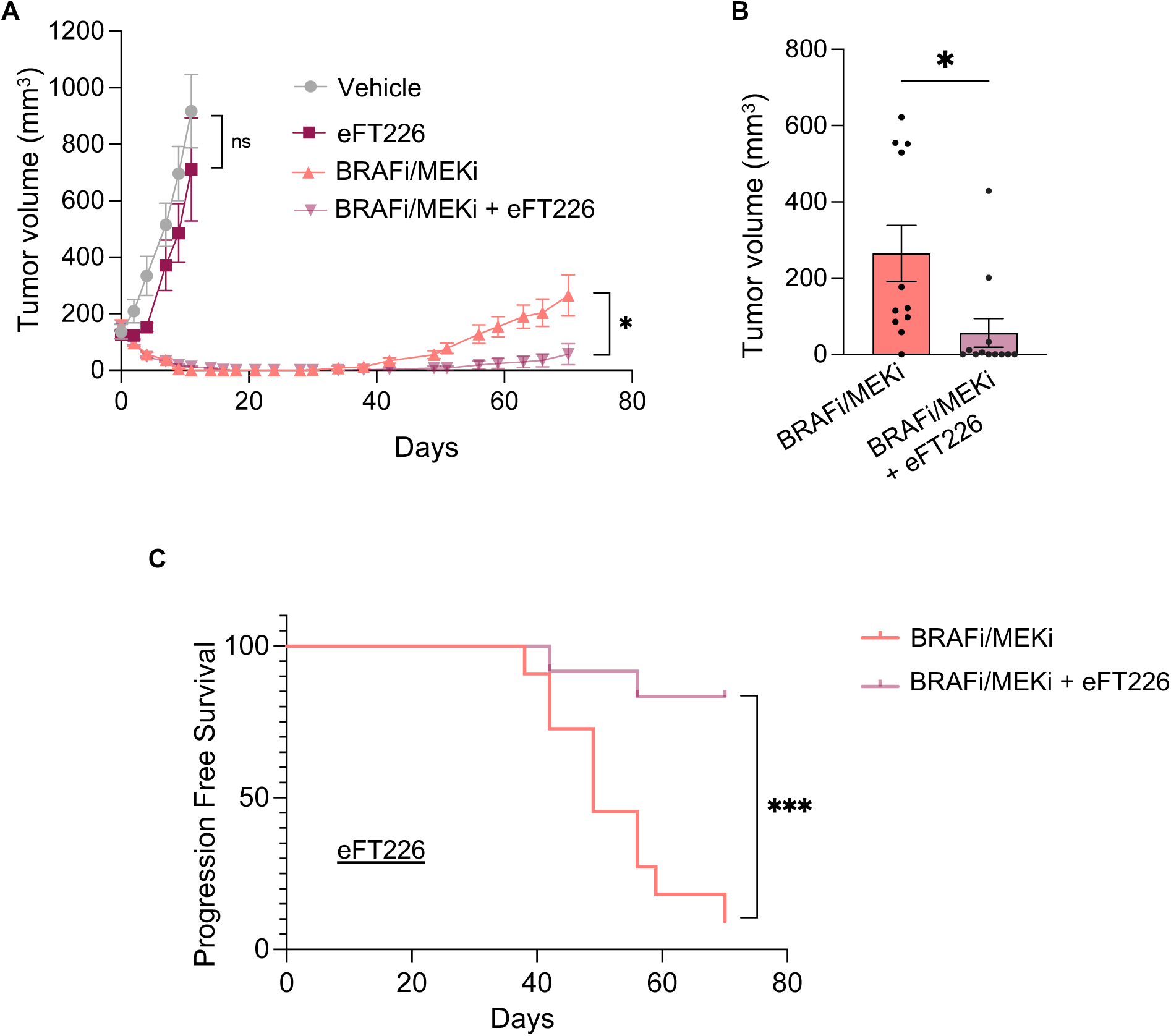
Targeting eIF4A with the use of eFT226 delays acquired resistance *in vivo*. **A**, Growth curve illustrating the changes in tumor volume over 70 days in a human BRAF-mutant A375 melanoma xenograft model treated with single and combined agents as indicated (Vehicle, n = 5; eFT226, n=5; BRAFi/MEKi, n = 11; BRAFi/MEKi + eFT226, n=12). Graphs represent mean values ± SEM. p-value was calculated by unpaired, two-tailed Student’s t-test (* p ≤ 0.05). **B**, Graphs representing the tumour volume (mean ± SEM) after 70 days of BRAFi/MEKi treatment (n=11) or with BRAFi/MEKi + eFT226 (n=12). p-value was calculated by unpaired, two-tailed Student’s t-test (* p ≤ 0.05). **C**, Kaplan-Meier curve for A375 melanoma xenograft treated with BRAFi/MEKi (n = 11), and BRAFi/MEKi + eFT226 (n=12). Median time progression was 49 days for BRAFi/MEKi group, and only 20% of mice belonging to the combination group relapsed after 70 days of treatment. Log rank (Mantel-Cox) for combination versus BRAFi/MEKi: p = 0.0004 (∗∗∗) and hazard ratio (log rank): 0.118 (95%, CI, 0.03-0.3).

In addition to eFT226, we chose to evaluate a series of synthetic flavagline derivatives for their capacity to inhibit the translation of luciferase reporter mRNA containing polypurine (AG)_10_ motif repeats within the 5’UTR, using an in vitro rabbit reticulocyte lysate system (Fig. S7, Supplementary Information). Among the compounds screened, RBX09 exhibited the strongest inhibitory activity (Fig. S7). RBX09 was generated by introducing an oxazolidinethione moiety fused to the flavagline scaffold (Supplementary Information). Given the promising activity of racemic RBX09, we prepared its pure enantiomer RBX0901 (Supplementary Information), which demonstrated greater effects than eFT226, with an approximately 100-fold improvement in EC₅₀, in inhibiting the translation of mRNA reporters containing AG-rich 5′UTRs compared to reporters with (UC)-rich or (AC)-rich or (UG)-rich 5′UTRs (Fig. 7A-C). As for eFT226, RBX0901 efficiently decreased the translation of the *53BP1* 5’UTR containing reporter in drug-tolerant cells, with no effect on the translation of the Renilla reporter containing no 5’UTR “Empty” (Fig. 7D). Additionally, RBX0901 tended to be more effective than eFT226 in reducing the translation of reporters containing the 5′UTRs of c-Myc and Cyclin D1, whose translation have previously been shown to be eIF4F-dependent (42,43), while having no impact on the translation of the reporter containing the 5’UTR of the housekeeping gene Tubulin (Fig. S8A). In line with this, RBX0901 had markedly reduced Cyclin D1 protein levels in A375 cells (Fig. S8B). Assessment of cytostatic activity revealed that both compounds exhibited similar efficacy in inhibiting A375 cell proliferation, resulting in comparable IC₅₀ values (eFT226: 3.7 nM; RBX09: 2.6 nM) (Fig. S8C). Combining BRAFi/MEKi with RBX0901, administrated at a non-toxic dose of 1mg/Kg twice a week (as for eFT226) (Fig. S9), significantly delayed tumor relapse, with 9 out of 14 mice remaining relapse-free (Fig. 7E,F). These findings demonstrate that co-treatment with BRAFi/MEKi and eIF4A inhibitors can effectively delay tumor relapse. Consistently, combining BRAFi/MEKi targeted therapy with eFT226 or RBX0901 also markedly reduced the emergence of drug-tolerant-derived proliferative clones *in vitro* (Fig. 7G-J).

**Figure 7:**
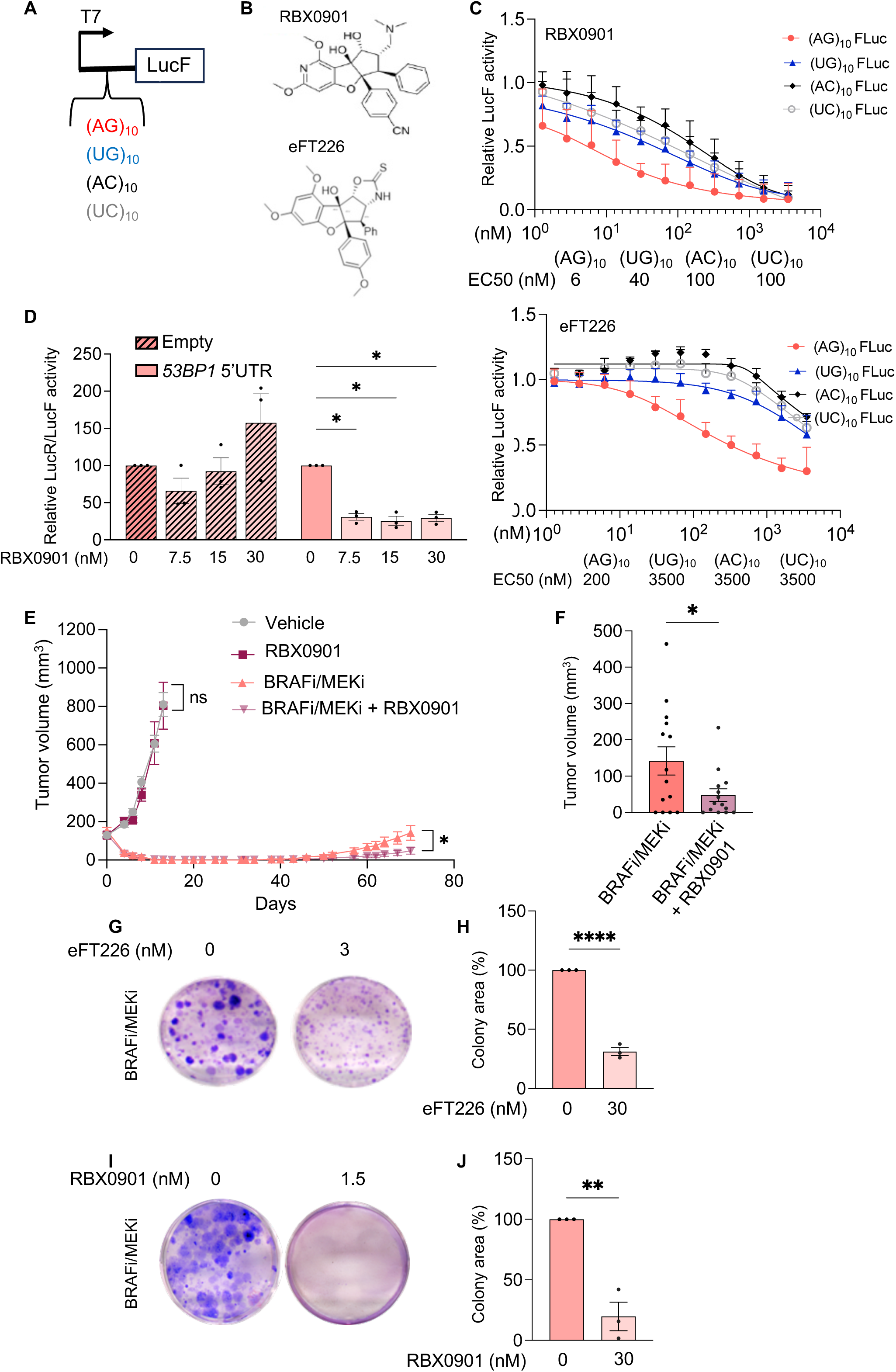
RBX0901, a novel flavagline derivative, exhibits comparable efficacy to eFT226 in delaying acquired resistance. **A**, Schematic of the luciferase reporters containing 5’UTRs with 10-mer sequence motif repeats used for *in vitro* translation. **B**, Chemical structures of RBX0901 and eFT226. **C**, Translation of luciferase mRNA reporter in rabbit reticulocyte lysate treated with increasing concentrations of eFT226 or RBX0901 for 1.5h. EC50 (nM) for each reporter is indicated. Data represent the mean ± SEM of 4 independent experiments. **D**, Quantification of the luciferase assay performed with 53BP1 5’UTR or Empty (without 53BP1 5’UTR) reporters in A375 drug-tolerant cells treated with the indicated concentrations of RBX0901 for 12 hours. Renilla (LucR) activity was measured at the end of drug treatment and the activity of the Firefly luciferase (LucF) was used as control of transfection. Data represent the effect of RBX0901 on the luciferase activity of the reporter containing 53BP1 5’UTR or the Empty vector and are normalized to the RBX0901 untreated conditions. The mean ± SEM from 3 independent experiments is reported (* p ≤ 0.05). **E**, Growth curve illustrating the changes in tumor volume over 70 days in a human BRAF-mutant A375 melanoma xenograft model treated with single and combined agents as indicated (Vehicle, n = 6; RBX0901, n=6; BRAFi/MEKi, n = 14; BRAFi/MEKi + RBX0901, n=14). Graphs represent mean values ± SEM. p-value was calculated by unpaired, two-tailed Student’s t-test (* p ≤ 0.05). **F**, Graphs representing the tumour volume (mean ± SEM) after 70 days of BRAFi/MEKi treatment (n=14) or with BRAFi/MEKi + RBX0901 (n=14). p-value was calculated by unpaired, two-tailed Student’s t-test (* p ≤ 0.05). **G**, Colony assay performed 3 weeks after incubating A375 cells drug tolerant cell (10000 cells) with BRAFi/MEKi (100 nM/ 10 nM) in the presence or absence of eFT226 (3 nM). **H**, Quantifications of the colony assay in G (mean ± SEM) from 3 independent experiments. p-values were calculated by unpaired, two-tailed Student’s t-test (**** p ≤ 0.0001). **I**, Colony assay performed 3 weeks after incubating A375 cells drug tolerant cell (10000 cells) with BRAFi/MEKi (100 nM/ 10 nM) in the presence or absence of RBX0901 (1.5 nM). **J**, Quantifications of the colony assay in I (mean ± SEM) from 3 independent experiments. p-values were calculated by unpaired, two-tailed Student’s t-test (** p ≤ 0.01).

## DISCUSSION

Here we report that eIF4A-mediated translation of the mRNA encoding the key error-prone NHEJ mediator 53BP1 mediates drug-induced mutability, driving melanoma cells evolution towards acquired resistance. Our findings reveal how the dynamic translation regulation of gene expression in response to DNA-damaging targeted therapy may facilitate melanoma cells adaptation to the DNA insult, while concomitantly and inadvertently enhance the mutation rate of cancer cells through the promotion of NHEJ pathway.

NHEJ pathway of DNA repair has been described as a survival strategy exploited by various cancers to withstand radio- or chemotherapy (44–47), with increased levels of key NHEJ components, including 53BP1, being correlated with therapeutic failure and cancer progression (48–51). NHEJ can compensate for defects in HDR (15), a vulnerability that underpins the clinical efficacy of poly(ADP-ribose) polymerase (PARP) inhibitors in the treatment of BRCA1/2-mutated cancers (52,53). Our findings, along with those of others, indicate that treatment with BRAFi/MEKi induces a BRCA-ness phenotype, characterized by the downregulation of key HDR components and a resulting impairment in HDR proficiency. Notably, in melanoma and other cancers subjected to targeted therapies, this repression has been shown to occur at the transcriptional level, at least for specific HDR factors such as RAD51 and BRCA1/2, and to be independent of cell cycle distribution (9,10,14,54). Nonetheless, a recent study identified RNA-binding proteins (RBPs) as contributors to HDR deficiency across multiple cancers (55), highlighting a possible role of post-transcriptional mechanisms in inhibiting the expression of HDR components, which remains yet to be tested.

In the context of HDR defects, our results demonstrate that drug-tolerant melanoma cells become increasingly reliant on the NHEJ pathway for DNA repair, a shift associated with elevated mutation frequency. While we used the *HPRT* locus as a proxy for global genomic mutability, our findings are consistent with recent results in colorectal cancer, where mathematical modeling and a modified Luria–Delbrück fluctuation assay indicated that targeted therapies are associated with an increased mutation rate (56). This shift away from high-fidelity DNA repair has been proposed to contribute to the genomic instability observed in cancer cells exposed to non-genotoxic treatments (10). While we cannot exclude the contribution of HDR defects to the observed mutation rate, our data support prior studies implicating NHEJ activity in genomic instability and acquired resistance of melanoma to targeted therapies (16).

Beyond transcriptional regulation, our study connects therapy-induced translational reprogramming to NHEJ efficiency and therapy-induced mutability. We identify eIF4A as a key regulator of *53BP1* mRNA translation, implicating the eIF4F complex in this process. Supporting this, *53BP1* mRNA translation was previously shown to be regulated by eIF4G1 in breast cancer cells following DNA damage (57)⍰, further confirming its reliance on eIF4F complex components. The translational control of 53BP1 by the eIF4F complex may thus be conserved across different cancer types and stress responses. Since the eIF4F complex is located at the convergence of several cell signaling pathway (58), including the PI(3)K/AKT/mTOR pathway and the RAS/RAF/MEK/ERK/MNK MAPK pathway, it is possible that the translation of the *53BP1* mRNA is regulated in cells tolerant to the various targeted therapies developed against these main signaling pathways in different cancer types. Nevertheless, eIF4F-independent functions of eIF4A have also been reported (59), and the precise mechanism by which eIF4A regulates *53BP1* translation remains to be elucidated. Future studies aimed at identifying additional trans-acting factors that cooperate with eIF4A in driving 5′UTR-dependent translation of *53BP1* will help clarify this mechanism.

While eIF4A inhibition has pleiotropic effects, our use of 5’UTR deletion provides evidence that the 53BP1-NHEJ axis is a critical mediator of the adaptive mutability of drug-tolerant cells, as shown in Figures 4 More specifically, experiments involving CRISPR/Cas9-mediated deletion of the 53BP1 5’UTR directly show that the translational upregulation of 53BP1 is not just correlated with, but is necessary and sufficient for, the observed increase in NHEJ activity. This provides a direct causal link between a specific translational event and the subsequent shift in DNA repair pathway choice. Indeed, although NHEJ, together with 53BP1 nuclear recruitment, is predominant during the G0/G1 phase of the cell cycle (13,27,60), and although BRAFi/MEKi targeted therapy has been shown to promote G0/G1 cell-cycle arrest in the majority of the cell population(61,62), our findings suggest that a direct mechanism of translational regulation, rather than an indirect effect of cell-cycle phase, is responsible for the observed shift in NHEJ. Moreover, this shift enhances genomic instability and the evolution towards acquired resistance. A detailed study will nevertheless be necessary to identify the *cis*-acting elements present in the 53BP1 5’UTR. We previously demonstrated that increased N6-methyladenosine modification in in the 5’UTR of a subgroup of mRNAs promoted enhanced translation in melanoma drug-tolerant cells (20), suggesting a possible mechanism for 53BP1 translational regulation.

Previous work has shown that the selective translation of DNA repair factors in response to genotoxic stress can be driven by structures or sequence elements within UTRs, often through cap-independent mechanisms (63). Although our previous genome-wide analysis of differentially translated mRNAs in this context did not identify other NHEJ factors, it will be important to investigate whether additional NHEJ proteins upregulated during targeted therapy are subject to similar translational regulation. Notably, LIG4 has recently been reported to undergo translational regulation following genotoxic stress (64).

Importantly, our data show that increased translation of *53BP1* alone is sufficient to drive the observed increase in mutation frequency in drug-tolerant cells. We propose and validate a therapeutic strategy to counteract this resistance mechanism by inhibiting eIF4A. Using two different small molecule inhibitors, including a newly developed one, we show that co-treatment with BRAF/MEK inhibitors significantly delays tumor relapse in xenograft models. The *in vivo* efficacy of eIF4A inhibition likely represents a composite effect derived from the translational suppression of numerous pro-survival pathways, and the precise contribution of the 53BP1-mutability axis relative to these other effects has not been quantitatively determined. However, DNA repair genes appear to exhibit eIF4A dependency in multiple cancer models (65,66), and the effect of eIF4A inhibition on NHEJ was phenocopied by selective targeting of 53BP1 mRNA translation, suggesting that this axis is, at least in part, a key mediator of the response.

Finally, previous work demonstrated that coordinated inhibition of HDR and NHEJ factors in melanoma leads to synthetic lethality triggered by unresolved DNA damage(14). In this work, we propose that inhibiting 53BP1-dependent NHEJ hinder the selection of new, fitter phenotypes driven by increased genetic instability that are drivers of acquired resistance to targeted therapy, pointing for a new role of mRNA translation in acquired mutability of drug-tolerant cells.

## METHODS

### Patient samples

Metastatic melanoma patients treated with dabrafenib or dabrafenib + trametinib at Gustave Roussy Cancer Campus (Villejuif, France) provided written informed consent for the collection of tissue samples for research. The transfer of clinical data from Gustave Roussy to Curie Institute was approved by Curie’s board: CRI Data [DATA230161]. Data from matched tumor samples from 3 patients at both baseline and after treatment were obtained and the study was approved by the Gustave Roussy Scientific Committee (IRB No. 2023-249). Patient clinical data are described in Table S1.

### Mice experiments

Six-week-old female athymic nude mice (Hsd:Athymic Nude-Foxn1nu, Mus Musculus, from Envigo) were subcutaneously inoculated with 2 million A375 cells. When tumors reached an average volume of 150 mm^3^, mice were fed with the control rodent diet or with diet supplemented with 200 ppm PLX4720 (BRAF inhibitor, Medchem Express) and 7 ppm PD0325901 (effective concentration of MEK inhibitor, Medchem Express). eFT226 (1 mg/kg, Medchem Express) or RBX0901 (1 mg/kg) were injected intraperitoneally two times per week for the entire duration of the experiment (70 days). EFT226 and RBX0901 were dissolved in DMSO to make a 0.1 mg/ml stock solution and then diluted in 20% 2-Hydroxypropyl)-β-cyclodextrin (HPCD, #128446-35-5, BLDpharm) solution just before injection. Tumor growth was monitored three times/week in two dimensions using a digital caliper. Tumor volumes were calculated with the ellipsoid volume formula L x ω2 x 0.5, where L is length and ω is width. Animals were allocated to experimental groups so that the groups had similar mean tumor volumes before treatment initiation. The investigator was not blinded to the group allocation or when assessing the outcome.

Experiments were performed in accordance with the CCAC guidelines and approved by the ethical committee of the “Plateforme d’évaluation Préclinique” of Gustave Roussy (#33216202109241653613).

### Cell culture and generation of drug tolerant cells

A375 melanoma cell lines were purchased from ATCC and were grown in high glucose (4.5 g/L) Dulbecco’s modified Eagle medium (Biowest, #L0101-500) supplemented with 10% FBS, 2 mM L-glutamine and 1% penicillin streptomycin. M249 cell lines were obtained from Thomas Graeber’s laboratory at department of Molecular and Medical Pharmacology, University of California, Los Angeles, USA and were grown in Roswell Park Memorial Institute 1640 (Biowest, #L0500) medium supplemented with 10% FBS, 2 mM L-glutamine and 1% penicillin streptomycin. All cells were grown in 5% CO2 humidified incubator at 37°C and regularly tested for mycoplasma and authenticated by short tandem repeat (STR) profiling. Drug tolerant cells were generated as previously described (20) through continuous treatment with BRAFi PLX4032 (500 nM) in combination with MEKi Cobimetinib (500 nM) for 72 h (A375) or PLX4032 (500 nM) in combination with 25 nM cobimetinib (M249). ∼70% of cells were killed and detached by the combination treatment. Surviving attached cells were trypsinized, washed once in PBS and re-plated in drug-free medium. the recovered surviving cells were maintained in drug-free conditions for a period of 9 days.

### Antibodies, siRNA, shRNA, RiboMap probes and other reagents

The antibodies, compounds, small interfering RNAs (siRNAs), short hairpin RNA (shRNA) and antisense oligonucleotides sequences, as well qPCR primers, used in this study are listed in Table S2.

### Lentiviral infection

The pLKO.1 vector-based shRNA lentiviral constructs were purchased from Sigma TRC Mission shRNA library. HEK293T cells were transfected with 10 μg of the lentiviral construct, 5 μg of VSVG (Addgene #14888) and 10 μg of psPax2 (Addgene #12260) using Lipofectamine 2000 (Invitrogen™, #11668019). All the constructs are listed in Table S2. After 48 hours, the supernatant was collected, centrifuged to remove cell debris and filtered through a 0.45 μm filter, and polybrene was added to the medium before transduction to target cells. The same procedure was repeated on the second day. The cells were then selected with puromycin (2 μg ml−1). Knockdown efficiency was confirmed by qPCR.

### Quantitative proteomic analysis

#### Sample preparation

Untreated and drug-tolerant A375 cells were lysed in Urea buffer (8M Urea, 50mM Ammonium bicarbonate). Cell lysates were sonicated and then incubated at room temperature for 10 minutes. Afterwards, the samples were centrifuged at 20,000 x g for 10 minutes. Supernatants were collected and protein extracts were quantified using the BCA Protein Assay Kit (Thermo FisherTM, #23225). 10 µg of total protein cell exctract was reduced by incubation with 5mM dithiothreitol (DTT) at 57°C for 30 min and then alkylated with 10 mM iodoacetamide for 30 min at room temperature in the dark. Trypsin/LysC (Promega) was added at 1:100 (w/w) enzyme to substrate. Digestion was performed overnight at 37°C. Samples were then loaded onto homemade C18 StageTips (AttractSPE Disk Bio C18-100.47.20, Affinisep) for desalting. Peptides were eluted using 40/60 CH3CN/H2O with 0.1% formic acid and vacuum concentrated to dryness. Peptides were reconstituted in 0.3% TFA before liquid chromatography–tandem mass spectrometry (LC–MS/MS) as described previously (67).

#### LC-MS/MS analysis

Liquid chromatography (LC) was performed with a Vanquish Neo nanoLC system (Thermo Scientific) coupled to an Orbitrap Astral mass spectrometer (MS), interfaced by a Nanospray Flex ion source (Thermo Scientific). Peptides were injected onto a C18 column (inner diameter 75 µm x 50 cm double nanoViper PepMap Neo, 2 μm, 100 Å, Thermo Scientific) regulated at a temperature of 50°C, and separated with a linear gradient from 100% buffer A (100% H2O in 0,1% formic acid) to 28% buffer B (100% CH3CN in 0,1% formic acid) at a flow rate of 300 nL/min over 104 min. The instrument was operated in data-independent acquisition (DIA) mode. MS full scans were recorded on the Orbitrap mass analyzer in centroid mode for ranges 380-980 m/z with a resolution of 240,000, a normalized automatic gain control (AGC) target set at 500% and a maximum injection time of 5 ms. The DIAs in the Astral analyzer were performed in centroid mode for a mass range of 380-980 m/z with windows width of 2 Da (without overlap), a maximum injection time of 3ms and a normalized AGC target of 500% after fragmentation using higher-energy collisional dissociation (25% normalized collision energy).

### Data processing

For identification, the data were searched against the Homo sapiens (UP000005640) UniProt database using Spectronaut (v19.7; Biognosys) by directDIA+ analysis using default search settings. Enzyme specificity was set to trypsin and a maximum of two missed cleavage were allowed. Carbamidomethyl was set as a fixed modification and N-terminal acetylation and oxidation of methionine as variable modifications. The resulting files were further processed using myProMS (v3.10. https://github.com/bioinfo-pf-curie/myproms (68))

For protein quantification, XICs from proteotypic peptides shared between compared conditions (TopN) with missed cleavages and carbamidomethylation were allowed. Median and scale normalization at peptide level was applied on the total signal to correct the XICs for each biological replicate (N = 3). To evaluate the statistical significance of the change in protein abundance, a linear model (adjusted on peptides and biological replicates) was performed, and a two sided T-test was apply on the fold change estimated by the model. The p-values were then adjusted using the Benjamini–Hochberg FDR procedure.The list of proteins from the quantitative proteomic analyses are reported in Table S3.

### GSEA analysis

Cluster profiler (69) and the fgsea library (70) were used to perform gene set enrichment on the quantitative proteomic ratio values (Table S3). Then, the gsea_go() function, using default parameters, was used to calculate the gene set enrichment. The enrichment plot was performed with the enrichplot package by using the enrichmentPlot function.

### Western blot

Cells were lysed in RIPA buffer (50 mM Tris-HCl pH 7.4, 100 mM NaCl, 1% Nonidet P-40, 0.1% SDS, 0.5% Sodium deoxycholate with protease inhibitor). The protein quantification was measured using the BCA Protein Assay Kit (Thermo Fisher™, #23225). For immunodetection, membranes were blocked in a buffer containing 0.1% Tween-20 (TBST) and supplemented with 5% powdered milk and exposed to appropriate antibodies.

### Quantitative real-time PCR analysis

Total RNA was extracted using TRIzol-chloroform and treated with DNase I (TURBO DNA-free; Invitrogen™, #AM1907) according to the manufacturer’s instructions. cDNA was synthesized using SuperScript III reverse transcriptase (Invitrogen™, #18080044). Quantitative real-time PCR was performed using Power SYBR Green PCR Master Mix (Thermo Scientific™, #4367559).

### Colony assay

Drug tolerant cells were seeded in 6 -well plates and treated with the indicated drug combinations for 3 weeks. Cells were washed once with PBS, stained with a 20% ethanol solution containing 0.5% crystal violet (Sigma Aldrich, #C0775) for 10 min, washed with PBS and destained in tap water. Images were analysed by ColonyArea plugin on Fiji software (71).

### CRISPR/Cas9 5’UTR deletion

CRISPR/Cas9 gene editing was performed using 2 fluorescent plasmid-based vectors (Vector builder) co-expressing hCas9 nuclease and sequence-specific sgRNA targeting 53BP1 5’UTR (Table S2). EGFP and mCherry were used as fluorescent markers for transfected cells.

A375 cells were transfected with 4 µg of each plasmid using Lipofectamine 2000 (Invitrogen™, #11668019) for 72h. Cells co-expressing GFP and mCherry were sorted by FACS and seeded into 96-well plates at one cell per well. Single-cell-derived clones were expanded, and gene editing efficiency was assessed by Sanger sequencing.

### Random plasmid integration assay

Random plasmid integration assay was performed as previously described (33) with some modifications. Briefly, untreated and drug tolerant A375 cells were seeded in 6-well plates and the following day were treated with 30 nM eFT226 or DMSO for 24 hours. At day 3, cells were transfected with 2-2.5 μg gel-purified BamHI-EcoRI-linearized pEGFP-C1 plasmid for 48 hours. When siRNA treatment was included in the experiment, two rounds of siRNA transfection (30 nM) with Lipofectamine RNAiMAX (Invitrogen™, #13778150) were performed on days 2 and 3, followed by transfection with the pEGFP-C1 plasmid. In experiments performed in A375 WT and D 5’UTR clones, untreated and drug tolerant A375 cells from each cell line were seeded in 6-well plates and the following day were transfected with pEGFP-C1 plasmid for 48 hours.

On day 5, cells were collected, counted, seeded, and grown in medium lacking or containing 0.5 mg/mL G418. Transfection efficiency of the linearized pEGFP-C1 was determined by flow cytometry. The cells were incubated at 37 °C to allow colony formation for 2 weeks by refreshing media every 3-4 days. The cells were then stained with a 20% ethanol solution containing 0.5% crystal violet (Sigma Aldrich, #C0775) and counted using ImageJ. Random plasmid integration events (number of G418-resistant colonies) were normalized by the plating efficiency (number of colonies without G418) and by transfection efficiency. pEGFP-C1 plasmid was a kind gift from van Attikum’s lab (Leiden University Medical Center, Leiden, The Netherlands).

### HPRT mutagenesis assay

HPRT assay was performed as previously described (34,35) with some modifications. Briefly, untreated and drug tolerant A375 cells were cultured in drug-free media for 6 days to ensure removal of any remaining endogenous HPRT activity. When applicable, cells were then treated with 30 nM eFT226 or DMSO for 24 hours and then plated onto a 150mm dish and treated with 20 µM 6-TG. Media with fresh 6-TG was changed every 3-4 days for 15-20 days until visible colonies appeared. Cells were also seeded at low density in drug-free media to determine plating efficiency. The cells were then stained with a 20% ethanol solution containing 0.5% crystal violet (Sigma Aldrich, #C0775) and colonies were counted using ImageJ.

The HPRT mutation frequency was calculated as the ratio of the number of HPRT−mutant colonies in 6-TG media to the number of surviving colonies plated in complete media to determine clonal efficiency.

### AHA-mediated RIBOsome isolation

mRNA enrichment in translating ribosomes was analyzed using AHARIBO RNA System kit (#AHA-RM12, Immagina Biotechnology). The assay was performed according to the manufacturer’s protocol.

The extracted RNA was quantified using Qubit (Thermo Scientific) and 200–500 ng was used for subsequent RT-qPCR experiments.

### RiboMap assay

RiboMap assay was performed as previously described (30). The Splints and *IQGAP1*-specific Padlocks and Primers probes were designed by the previous study (30). The *53BP1*-specific hybridization regions in Padlocks and Primers probes were designed using Picky 2.2 software. To visualize translating mRNAs, fluorescent probes complementary to the DNA amplicons were used instead of *in situ* sequencing. Probes were purchased from IDT and probes sequenses are listed in Table S2.

Images were acquired using a Leica SP8X inverted confocal laser scanning microscope (CLSM) equipped with a 63X oil immersion objective (NA = 1.4). Sequential excitation was performed using a laser diode at 405 nm and a white light laser (WLL) at 488 nm, 594 nm and 650 nm. Emission was detected using GaAsP hybrid photon detectors and photomultiplier tube (PMT), with detection windows set to 415-465nm for 405 nm excitation, 500–550 nm for 488 nm excitation, 605-670nm for 594 excitation and 660–710 nm for 650 nm excitation. During acquisition, an additional channel was added to highlight cells and facilitate their detection for quantification. The BrightR mode on the Hybrid Detector was used, with the detection window set to 575–660 nm for excitation at 488 nm. Image acquisition and system control were performed using LAS X software (Leica Microsystems). At least 30 cells per sample were imaged, and the images were processed with Fiji (72) by using the MIC-MAQ plugin (https://github.com/MultimodalImagingCenter/MIC-MAQ). 3D Z-stack images are converted to 2D images by applying a Z maximum intensity projection. Nuclei and cells are segmented from the DAPI and the additional fluorescence images using the Cellpose deep-learning network (73) with the Cyto3 model. For cell, images are first We created artificially an other channel were autofluorescence of the cytoplasm was used to highlight cells contour. Segmentation is performed using a diameter corresponding to the expected size of nuclei or cells, typically set to 100 pixels for nuclei and 300 pixels for cells. To facilitate spot detection, the background is removed using the subtract background function with a rolling ball radius of 30 pixels. During analysis with MIC-MAQ, nuclei and cells are exported as individual regions in ZIP files, with one ZIP file per image in the project. Subsequently, morphological and intensity parameters—including area, mean intensity, and spot counting using the Find Maxima function are measured for each cell in Alexa Fluor 488, 594, 647 channels.

### Immunofluorescence and multiplex immunostaining

For immunostaining, untreated and drug-tolerant cells were plated in 6-well plates with coverslips and fixed with 4% PFA at day 1 after drug removal. When applicable, 24 hours treatment with eFT226 (30 nM) or DMSO was performed before fixation. Cells were permeabilized in 0.5% PBS-Triton, washed with PBS, and then blocked for 1 hour at room temperature in PBS containing 0.1% Tween® 20 Detergent (Sigma Aldrich #P6585) and 5% bovine serum albumin (Sigma Aldrich #A9576). Primary and secondary antibodies were diluted in blocking solution. Primary antibodies were incubated for 1 hour to 3 hours at room temperature, depending on the antibody, and secondary antibody were incubated for 1 hour at room temperature. Cells were then washed using PBS with 0.1% Tween® 20 (Sigma Aldrich #P6585) and coverslips were mounted on glass slides using ProLong™ Diamond with DAPI (Invitrogen™ #P36971) mounting medium. Fluorescent images were acquired on a Leica widefield DM6000B upright microscope equipped with a 63x oil objective (N.A=1.4) and coupled with a Hamamatsu sCMOS ORCA Flash4.0 camera (pixel size: 6.5µm). Alexa Fluor 488 DAPI were detected using bloc filters (DAPI: ex BP405/60 - em BP 470/50; FITC: ex BP470/40 - em BP525/50). The system is driven by MetaMorph® (Molecular Devices) software. Images were analyzed using a semi-automatic macro on Fiji software. Nuclei containing ≥ 10 distinct foci were defined as foci-positive, and the percentage of positive nuclei was calculated as [(number of foci positive nuclei) / (number of nuclei scored)]* 100. A minimum of 100 nuclei per sample were scored, and data were collected from three biological replicates.

For multiplex immunostaining, untreated and drug-tolerant cells at day 1 of drug removal were collected, pelleted, and embedded in OCT compound. The samples were then flash-frozen in liquid nitrogen. Frozen blocks were sectioned using a cryostat, and tissue sections were mounted onto poly-L-lysine–coated coverslips Akoya Biosciences CODEX multiplex immunostaining was performed according to its manufacturer’s.

In brief, coverslips were fixed with acetone, rehydrated and then stained with a combination of DNA-barcoded primary antibodies, including in-house conjugated antibodies, washed and post-fixed in ice-cold methanol. They were mounted on a CODEX system for multiple cycle immunostaining and imaged using a Keyence microscope with CODEX instrument manager and Keyence software (BZ-X800 viewer). Secondary antibodies were fed to the instrument in a pre-prepared 96-well plate. In total, 5 cycles of immunostaining (including 2 blanks with only nuclear staining) were run, consisting of DAPI nuclear staining, Atto550-, Cy5- and Alexa Fluor 488 fluorophores. CODEX processor (v. 1.7.0.6 and 1.8.3.14) performed automated image registration, extended depth of focus, shading correction, and autofluorescence and background subtraction.

### Multiplex immunostaining data analysis

Images were automatically analyzed using QuPath software (74). All images (in qptiff format) were imported into a QuPath project, and nuclei were automatically segmented on DAPI-stained images using the Cellpose deep learning algorithm (73). The model employed was Cyto2 with a cell diameter set to 15 pixels and a fixed pixel size of 0.5. Cell boundaries were estimated by expanding the detected nuclei by a maximum of 15 pixels and limiting cell size relative to the nucleus with a factor of 5. Mean intensity values of various markers were calculated for each cell and exported as comma-separated values (CSV) files for further analysis.

To visualize the data, heat maps were generated using GraphPad Prism. Each row in the data matrix represents a single cell, with color-coding reflecting the relative intensity of the corresponding marker signal. In the heat maps, cells were organized in ascending order based on 53BP1 expression levels.

### Immunohistochemistry (IHC)

IHC was performed using the BenchMark Ultra automated staining system (Roche Diagnostics). Antigen retrieval was carried out with the CC1 buffer (pH 8.0) for 64 minutes at 95°C. Tissue sections were then incubated with the anti-53BP1 antibody (Cell Signaling Technology, #88439) at 1:400 dilution for 1 hour at room temperature. Detection was performed using the UltraView Universal DAB Detection Kit (Roche) according to the manufacturer’s instructions.

### *In vitro* transcription

T7-LucF sequences were amplified by PCR using the pCREL plasmid as a template (75,76). The primers used are listed in Table S2. (AG)_10_-, (UC)_10_-, (AC)_10_- and (UC)_10_ reporter transcripts were synthesized using the *mMESSAGE mMACHINE™* T7 Transcription Kit (Invitrogen™, Cat. #AM1344) according to the manufacturer’s instructions. 0.2 µg of T7-LucF construct was used as the transcription template. The in vitro transcription reaction was carried out at 37 °C for 2 hours. Following transcription, template DNA was removed by digestion with TURBO DNase (Invitrogen™, #AM2238) and RNA was purified using phenol–chloroform.

### *In vitro* translation

*In vitro* translation assays were performed with Rabbit Reticulocyte Lysate System (Promega Cat. No. L4960) adding 0.5 ng of reporter mRNAs according to the manufacturer’s instructions. The drugs at the indicated concentrations were added to the reaction a for 1h30 min at 30°C prior to the measurement of luciferase activity using Dual-Luciferase Reporter Assay System (Promega, #E1910).

### Luciferase assay

Untreated, drug-tolerant cells and cells released from the drugs for 9 days were transiently transfected with psiCHECK ™-2 Vector (Promega, #C8021,) using Lipofectamine LTX Reagent (Invitrogen™, #15338100). In psiCHECK ™-2 Vector, 317 bp downstream the SV40 early enhancer/promoter transcription start site, corresponding to nt 372 to 684, were removed. 53BP1, Tubulin, c-Myc and Cyclin D1 5’UTR sequences were cloned at the level of NheI site, upstream to Renilla coding sequence. The *Firefly* luciferase signal served as an internal transfection normalization control. After 48 hours, part of the transfected cells was harvested for RNA extraction to measure RNA expression by RT-qPCR, while the rest of the cells were used to measure the luciferase activity using the Dual-Luciferase Reporter Assay System (Promega, #E1910). Data are presented as the ratio between Renilla and Firefly luciferase RNA expression or luminescence activities, respectively.

For experiments performed in the presence of drugs, treatments at the indicated concentrations were performed 24 hours after psiCHECK ™-2 vector transfection, for 12 (53BP1 5’UTR reporter) or 24 hours (Tubulin, c-Myc and Cyclin D1 reporters). At the end of drug treatment, luciferase activity was analyzed.

### Proliferation assay

A375 cells were seeded in triplicates in 96-well plates. The day after, cells were treated with the indicated concentrations of drugs. Live-cell proliferation was monitored using the IncuCyte S3 System (Sartorius). Images of each well were captured every 3 hours over a 72-hour period. Proliferation data were quantified based on confluency measurements, and values were normalized to the confluency at time zero (T0) to account for initial seeding variability.

For long term proliferation assay with Δ5’UTR clones, drug-tolerant cells were seeded into T25 flasks and treated the following day with 100 nM BRAFi and 10 nM MEKi. Images of each flask were acquired weekly for 8 weeks under continuous BRAFi/MEKi treatment using the IncuCyte® S3 Live-Cell Analysis System (Sartorius). Proliferation was quantified as percent confluence (plate coverage), and values were normalized to the confluence at time zero (T0) to account for variability in initial cell seeding. Confluence measurements were obtained using the image analysis module in the IncuCyte system.

### Quantification and statistical analysis

Statistical analyses were performed on GraphPad Prism (Version 10.3.0) and statistical significance defined as a P value < 0.05 was determined by two-tailed unpaired Student’s t test or, when indicated in the figure legends, by two-tailed unpaired Student’s t test. Comparisons in multiple groups were analyzed with one-way or two-way ANOVA unless otherwise indicated in the figures’ legends.

## ACKNOWLEDGEMENTS

This work was supported by grants from Institut Curie, Gustave Roussy, INSERM, CNRS, Equipe labellisée Ligue Nationale Contre le Cancer (LNCC), Cancéropole île de France, Université Paris-Saclay. LF was supported by post-doctoral fellowships from Fondation ARC pour la recherche sur le cancer, Fondation de France. Proteomics was performed by the CurieCoreTech Mass Spectrometry Proteomics (LSMP) supported by grants from La Région Ile de France (NEX061034) and ITMO Cancer of Alliance Nationale pour les Sciences de la Vie et de la Santé (Aviesan) and Institut National du Cancer (INCa) on funds administered by INSERM (21CQ016-00) for MS analysis. The LSMP thanks Patrick Poullet from the bioinformatics platform of the Institut Curie U1331 for the continuous development of myProMS and Michael Richard from the LSMP for statistical help.

The authors acknowledge the Light Microscopy facility from the Multimodal Imaging Center of the Institut Curie (CNRS UAR2016/ Inserm US43/ Institut Curie / Université Paris-Saclay). Data management, quality control and primary analysis were performed by the Bioinformatics platform of the Institut Curie.

## AUTHOR INFORMATION

Investigation and methodology: LF,LL, EG, HL, DB. Proteomics was performed by BL and DL. LF, LL, EG, CR and SV conceived the experiments and analyzed data. SV coordinated the whole study. LF wrote the initial version of the manuscript, SV and CR helped in writing the manuscript. Funding acquisition: SV, CR, LF.

## COMPETING INTERESTS

Authors declare that they have no competing interest.

## DATA AVAILABILITY

All data are available in the main text or the supplementary materials.

## Figure Legends

**Supplementary Figure S1:**
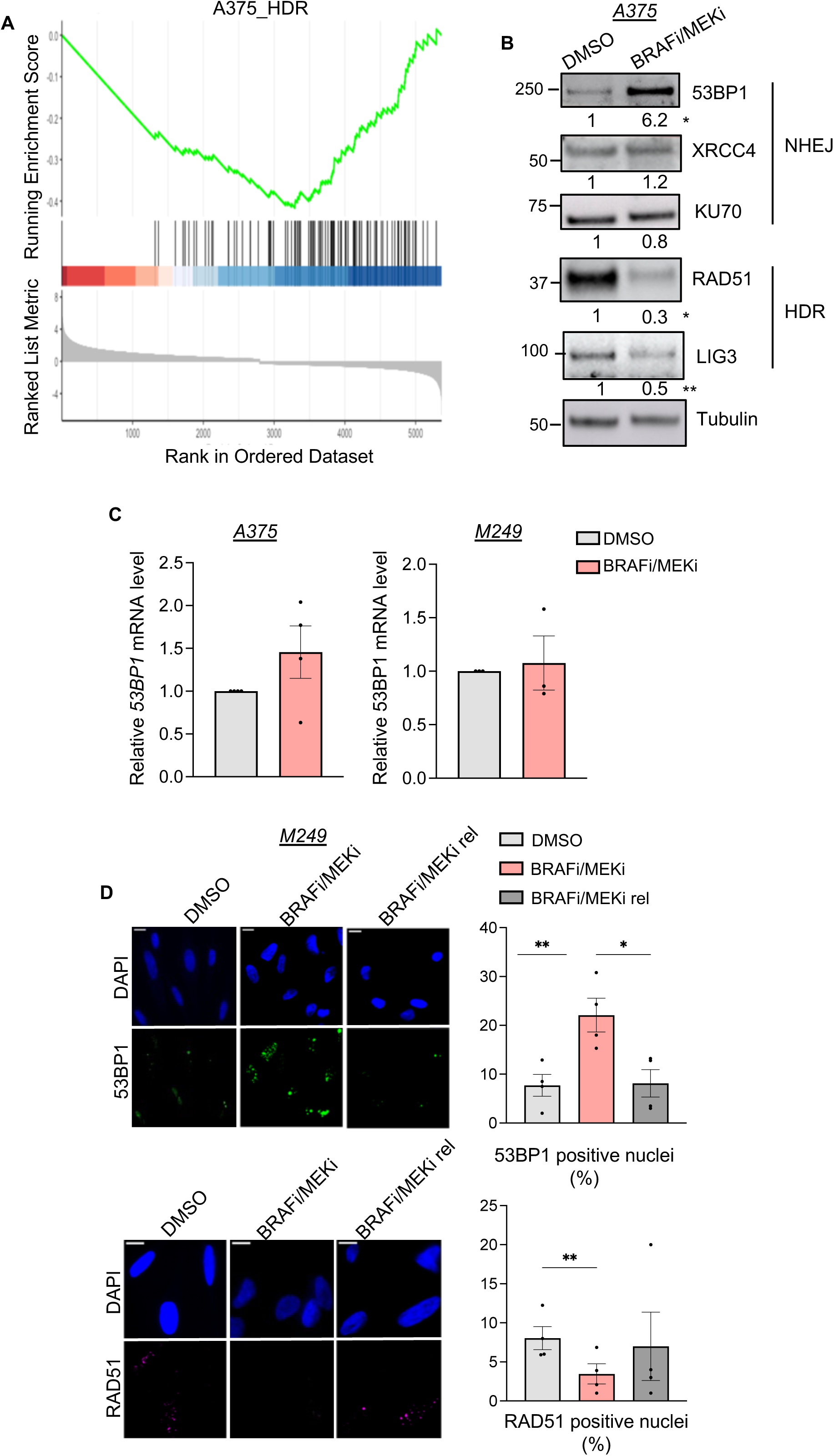
**A**, GSEA enrichment of the differentially expressed proreins. NES = −2.321, q value = 2.81e^-05^. **B**, Western blot illustrating 53BP1, XRCC4, KU70, RAD51 and LIG3 protein levels in A375 cells after 3 days of treatment with DMSO (DMSO) or BRAFi/MEKi treatment (BRAFi/MEKi). GAPDH serves as a loading control, and the relative quantification from 3 independent experiments is indicated. p-values were calculated by unpaired, two-tailed Student’s t-test (* p ≤ 0.05, ** p ≤ 0.01). **C**, RT-qPCR quantification of *53BP1* mRNA level in control cells (DMSO) or in BRAFi/MEKi drug tolerant cells. Data represent the mean ± SEM from 4 (A375 cells) or 3 (M249 cells) independent experiments. **D**, Representative images of immunofluorescence in M249 control (DMSO), drug tolerant cells (BRAFi/MEKi) and drug tolerant cells cells released from targeted therapy for 9 days (BRAFi/MEKi rel) stained for 53BP1 (green) and nuclei (DAPI, blue). (bottom) Representative images of immunofluorescence in cells stained for RAD51 (magenta) and nuclei (DAPI, blue). Scale bar, 10 μm. The quantification of the signal analyzed by percentage of cells with ≥ 10 foci/nucleus is reported and data represent the mean ± SEM from 4 independent experiments. p-values were calculated by paired, two-tailed Student’s t-test (* p ≤ 0.05, ** p ≤ 0.01).

**Supplementary Figure S2:**
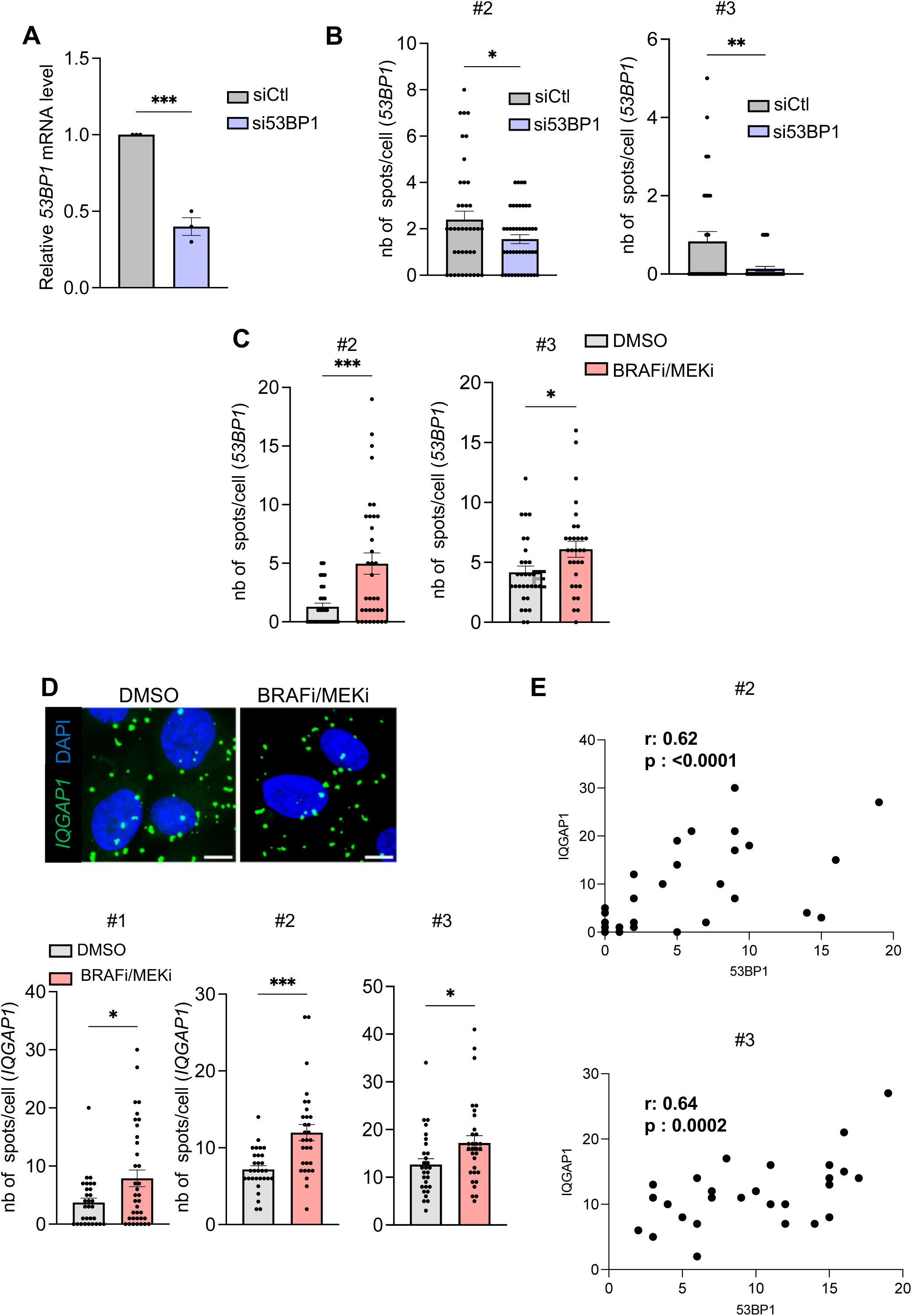
**A**, RT-qPCR quantification of *53BP1* mRNA level in A375 cells transfected with control siRNA (siCtl) or with siRNA targeting *53BP1* (si53BP1) corresponding to Fig. 2D,E. Data represent the mean ± SEM from 3 independent experiments (*** p ≤ 0.001). **B**, Quantification of translating *53BP1* mRNAs (spots) in A375 cell transfected with siCtl or si53BP1. Data shown represent the mean ± SEM of the number of spots/cells quantified in other two independent experiments (corresponsing to Fig. 2E) (* p ≤ 0.05, ** p ≤ 0.01). **C**, Quantification of translating *53BP1* mRNAs (spots) in A375 control cells (DMSO) or in cells surviving targeted therapy (BRAFi/MEKi). Data shown represent the mean ± SEM of the number of spots/cells quantified in other two independent experiments (corresponsing to Fig. 2G) (* p ≤ 0.05, *** p ≤ 0.001). **D**, (top): Representative confocal images of translating *IQGAP1* mRNAs (green) assessed by RiboMap assay in A375 control cells (DMSO) or in cells surviving targeted therapy (BRAFi/MEKi). Nuclei are stained with DAPI (blue). Scale bar: 10 µm., (bottom): Quantification of translating *IQGAP1* mRNAs (spots). Data represent the mean ± SEM of the number of spots/cells quantified in 3 independent experiments (* p ≤ 0.05, *** p ≤ 0.001). **E**, Scatterplot illustrating the correlation between translating *53BP1* mRNAs and *IQGAP1* mRNAs found in other two biological replicates in A375 drug tolerant cells (corresponding to Fig. 2I). Pearson correlation coefficient (r) and p-value are reported.

**Supplementary Figure S3:**
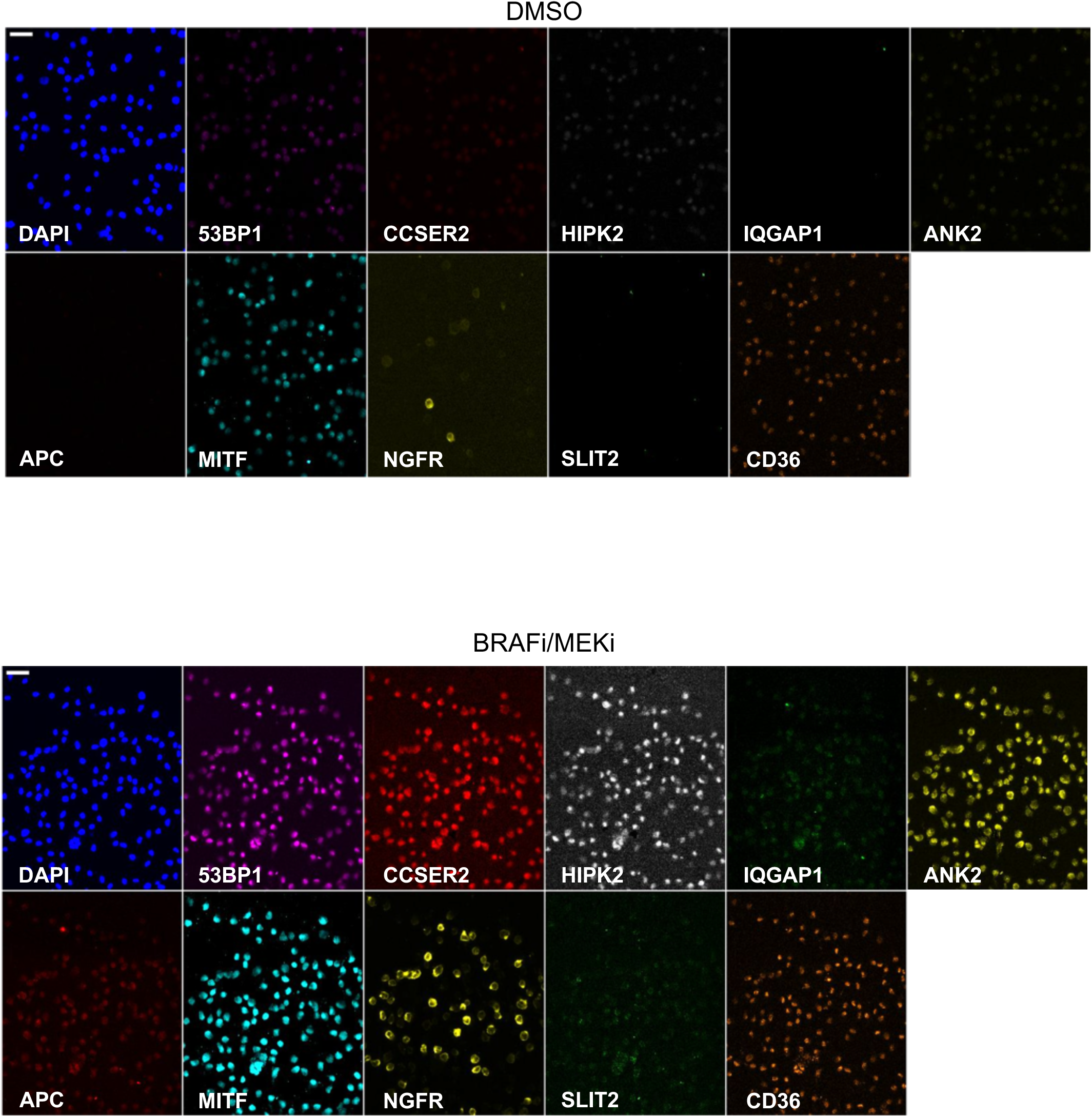
**A**, Quantification of *53BP1* mRNA enrichment in active ribosomes performed using RT-qPCR in A375 WT cells and in A375 cells deleted for 53BP1 5’UTR (Δ5’UTR, clone 1.11) as described in Figure 3. Control cells (DMSO) or drug tolerant cells that survived targeted therapy (BRAFi/MEKi) were analyzed for *53BP1* mRNA enrichment, and data represent the mean ± SEM from 3 independent experiments (** p ≤ 0.01, *** p ≤ 0.001). **B**, Representative images of immunofluorescence in A375 Δ5’UTR cells (clone 1.11) treated with DMSO (control) or in A375 Δ5’UTR cells that survived BRAFi/MEKi targeted therapy (BRAFi/MEKi) stained for 53BP1 (green) and nuclei (DAPI, blue). Scale bar, 10 μm. **C**, Quantification of 53BP1 signal in the conditions described in B, analyzed by percentage of cells with ≥ 10 foci/nucleus. Data represent the mean ± SEM from 4 independent experiments. **D**, RT-qPCR quantification of *53BP1* mRNA level in A375 cells stably expressing control shRNA, or shRNA targeting *53BP1*. Data represent the mean ± SEM from 3 independent experiments (*** p ≤ 0.001).

**Supplementary Figure S4:**
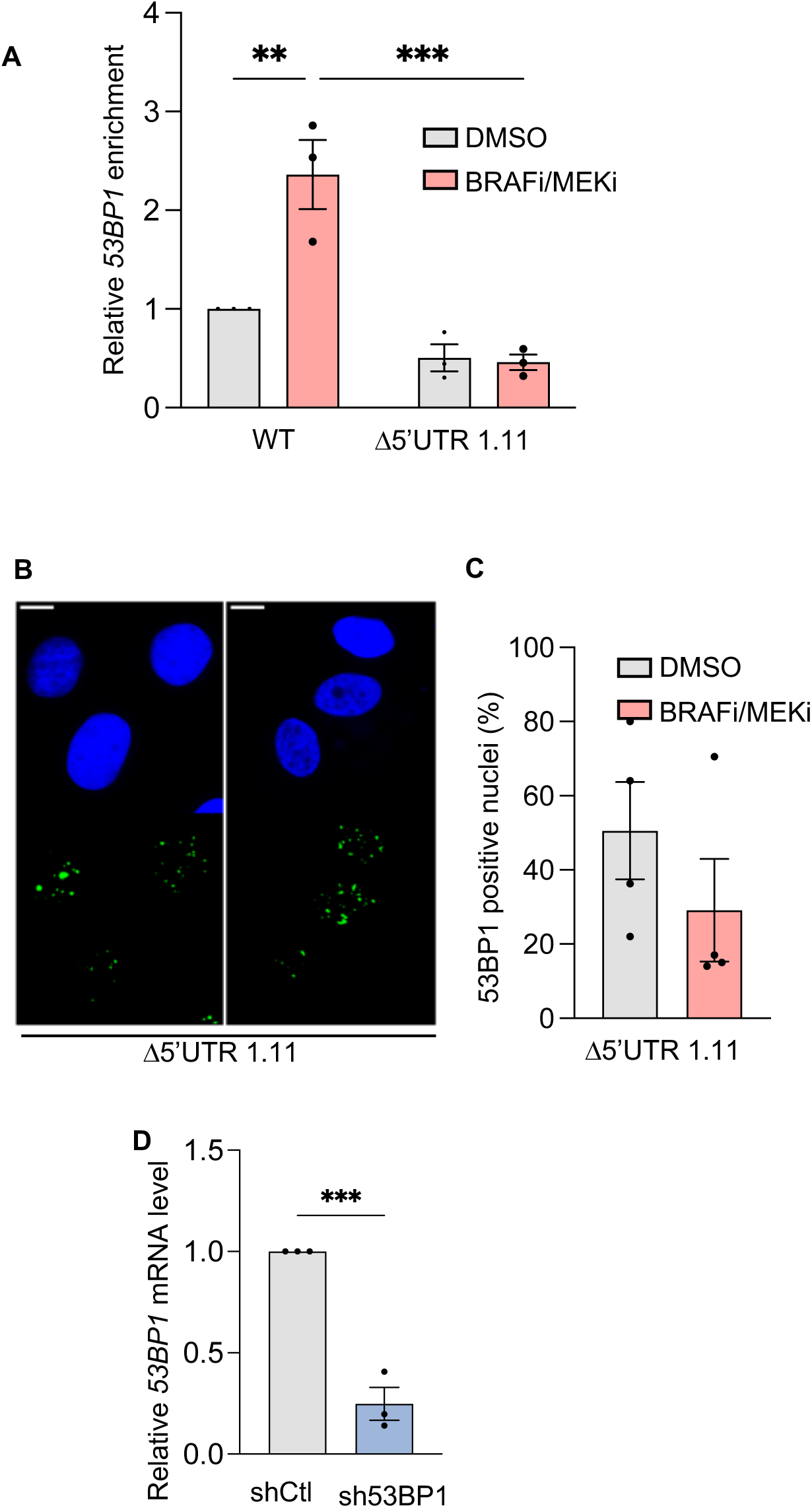
Representative images of translationally regulated (53BP1, CCSER2, HIPK2, IQGAP1, ANK2, APC) or transcriptionally regulated (MITF, NGFR, SLIT2, CD36) protein markers analyzed by multiplex immunostaining (CODEX) in A375 control cells (DMSO) or in cells surviving 3 days treatment with targeted therapy (BRAFi/MEKi).

**Supplementary Figure S5:**
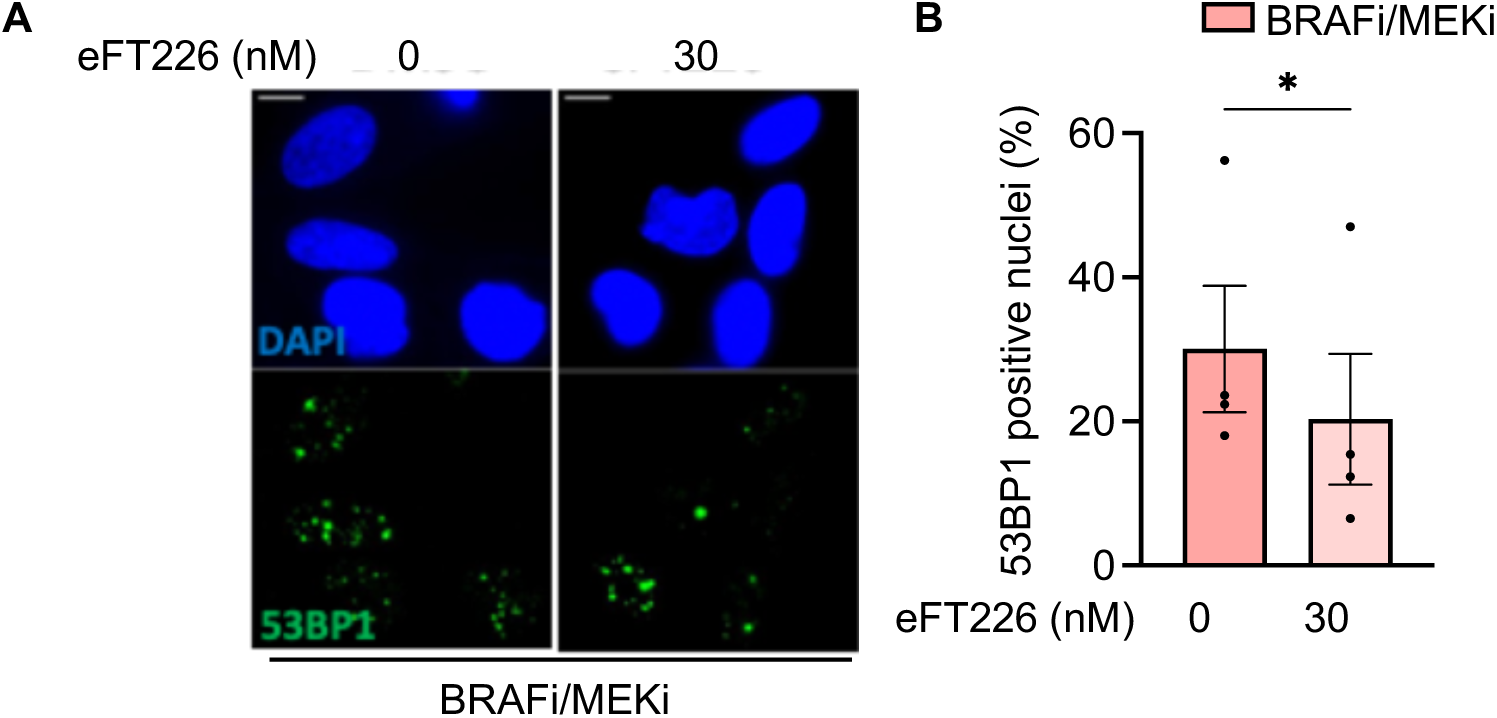
**A**, Representative images of immunofluorescence in A375 drug tolerant cells that survived BRAFi/MEKi targeted therapy (BRAFi/MEKi), treated with or without 30 nM eFT226 for 24 hours and stained for 53BP1 (green) and nuclei (DAPI, blue). Scale bar, 10 μm. **B**, Quantification of 53BP1 signal of the conditions in A, analyzed by percentage of cells with ≥ 10 foci/nucleus. Data represent the mean ± SEM from 4 independent experiments. p-values were calculated by paired, two-tailed Student’s t-test (* p ≤ 0.05).

**Supplementary Figure S6:**
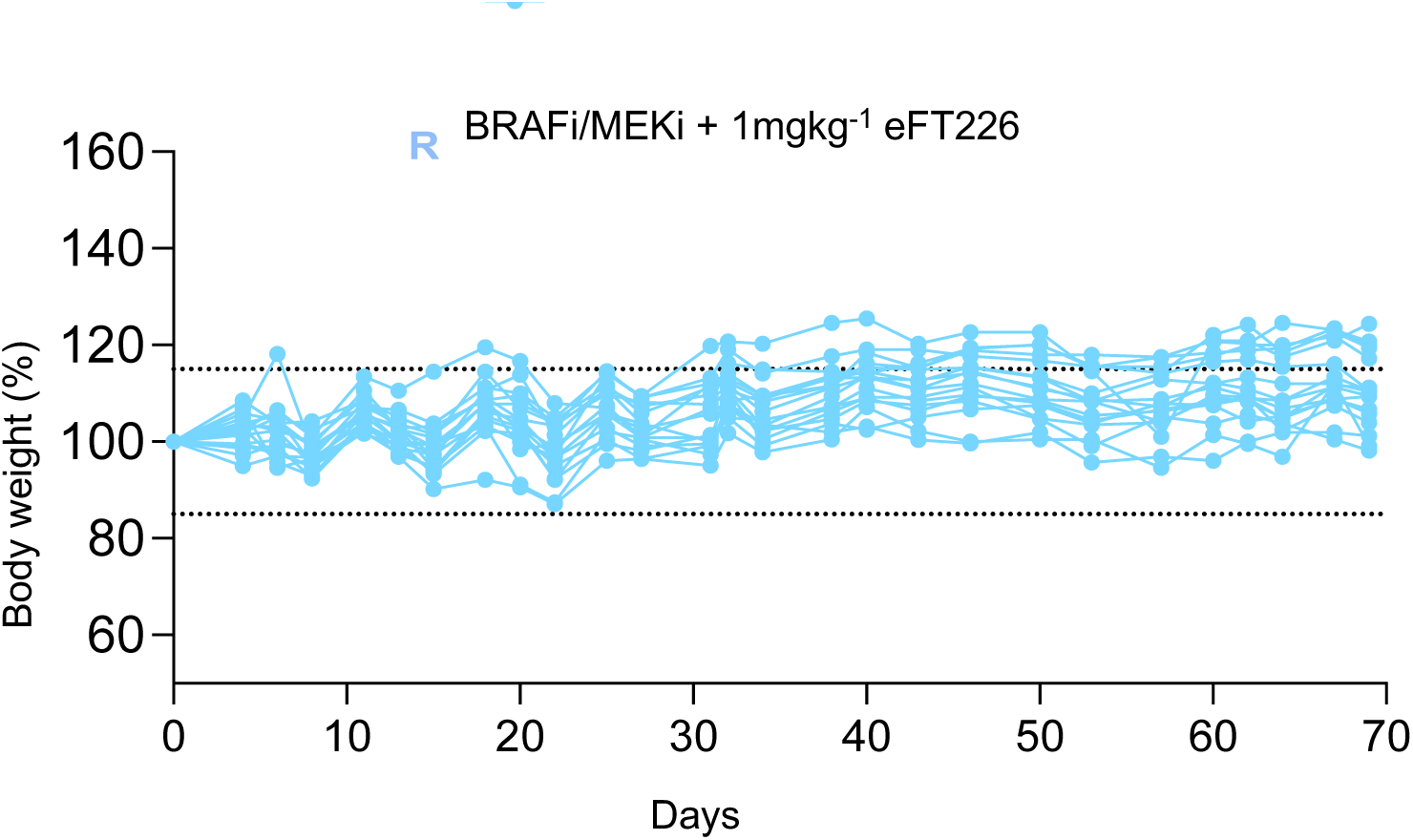
Graph showing the percentage variation in body weight of mice after 70 days of treatment with PLX4720 (BRAFi) and PD0325901 (MEKi), combined with eFT226 (1 mg/kg).

**Supplementary Figure S7:**
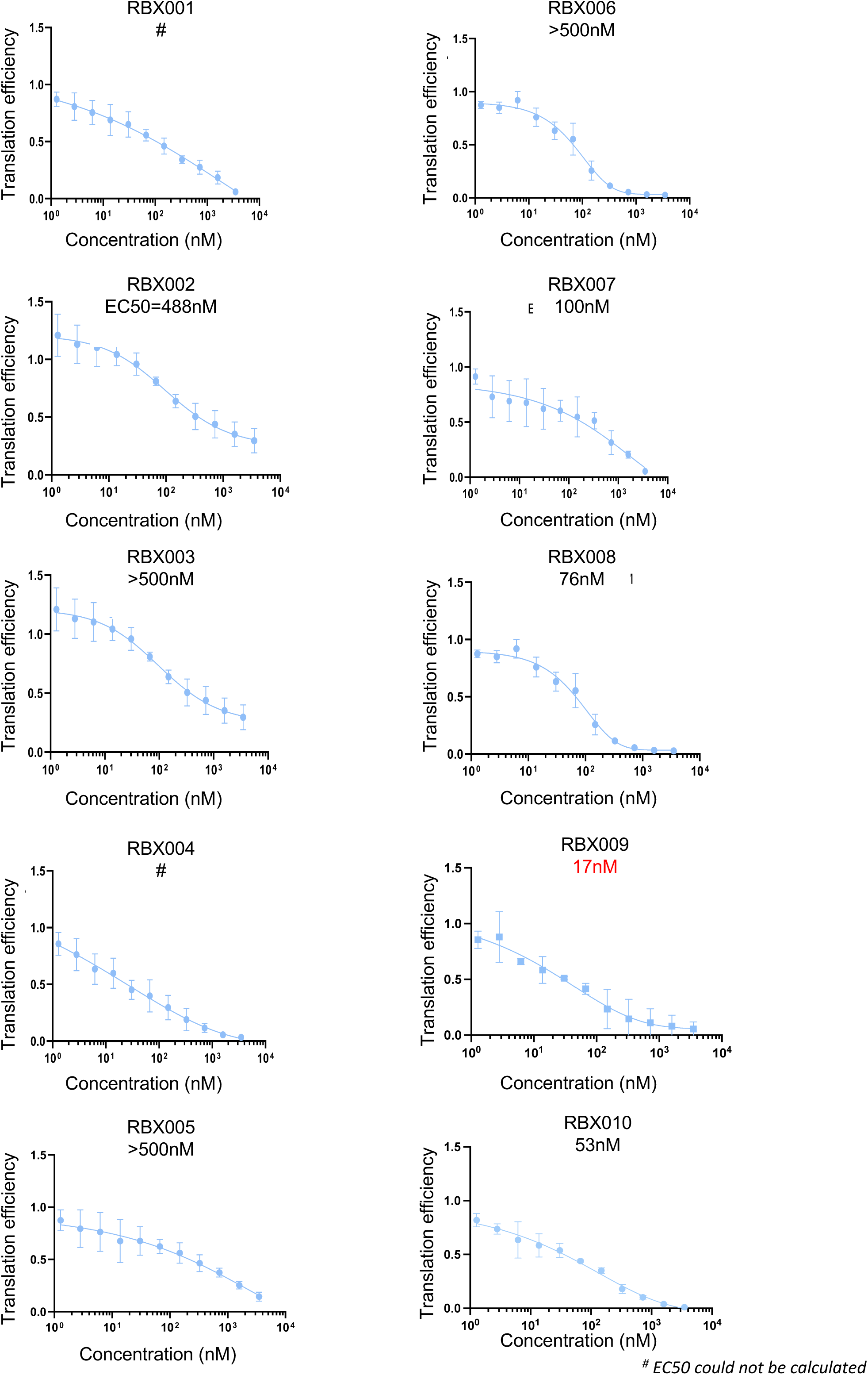
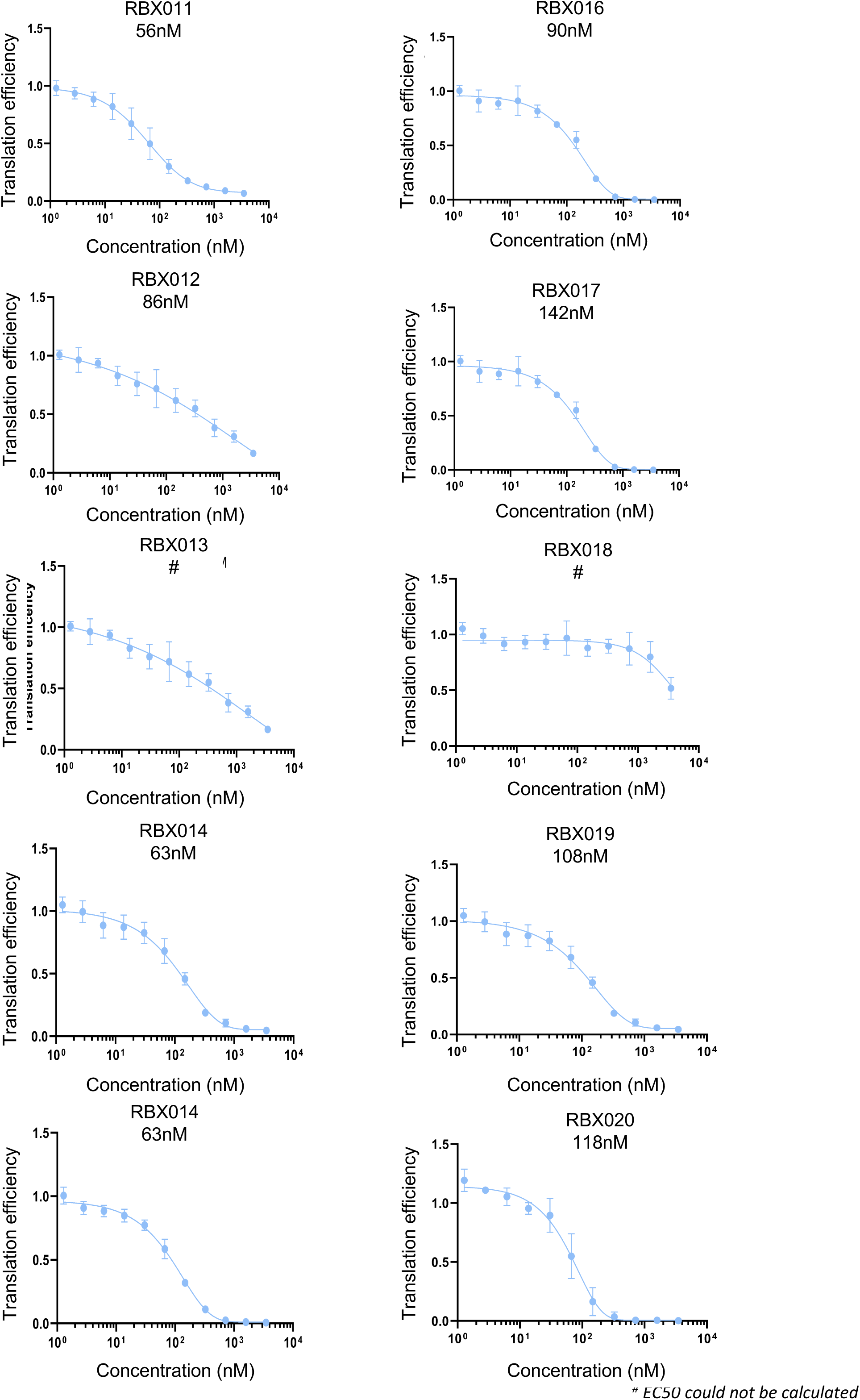
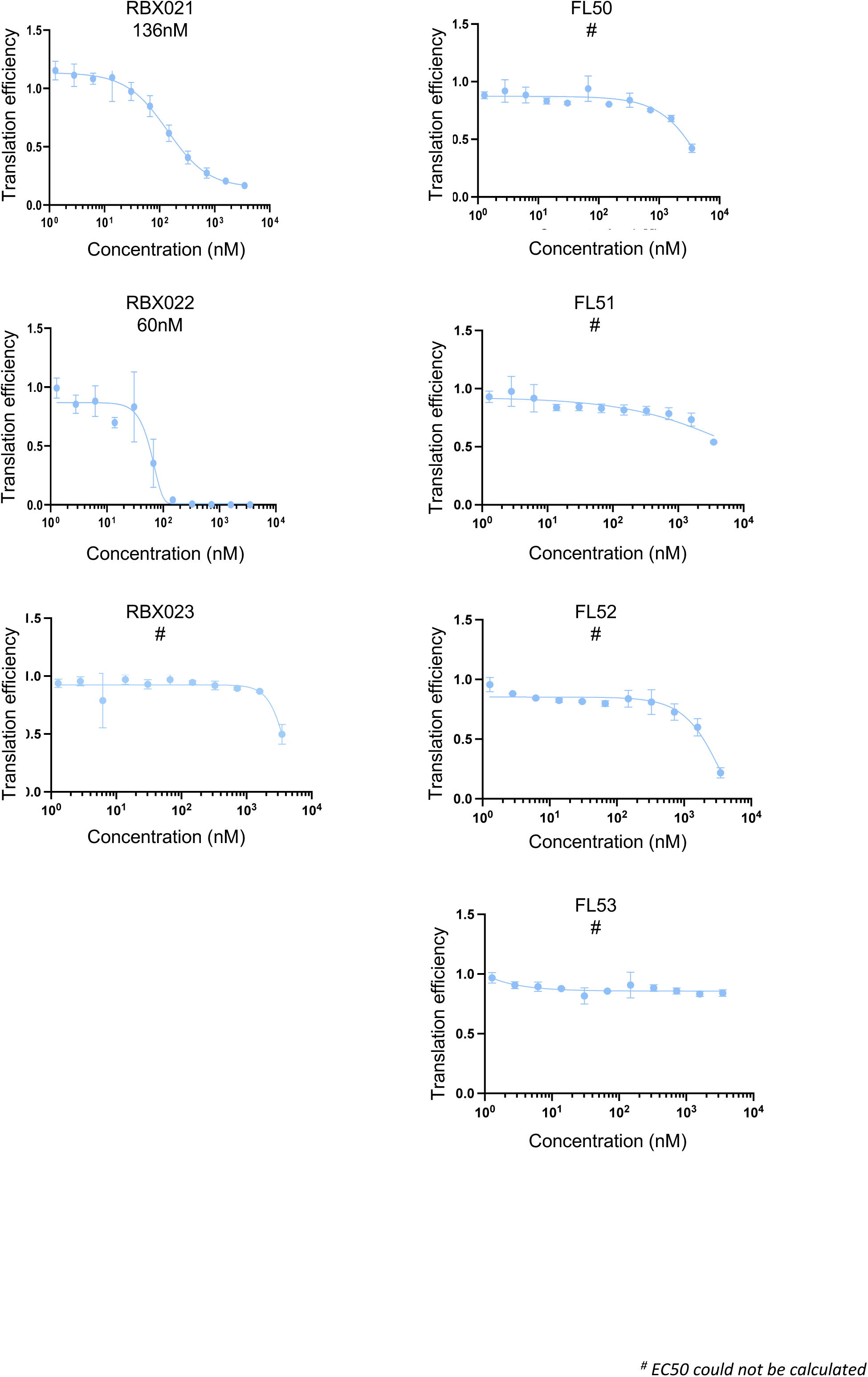
Screening of flavagline derivative compounds performed by assessing the in vitro translation of the (AG)₁₀ luciferase mRNA reporter in rabbit reticulocyte lysate treated with increasing concentrations of compounds for 1.5 hours. Where applicable, the EC₅₀ (in nM) is indicated for each compound.

**Supplementary Figure S8:**
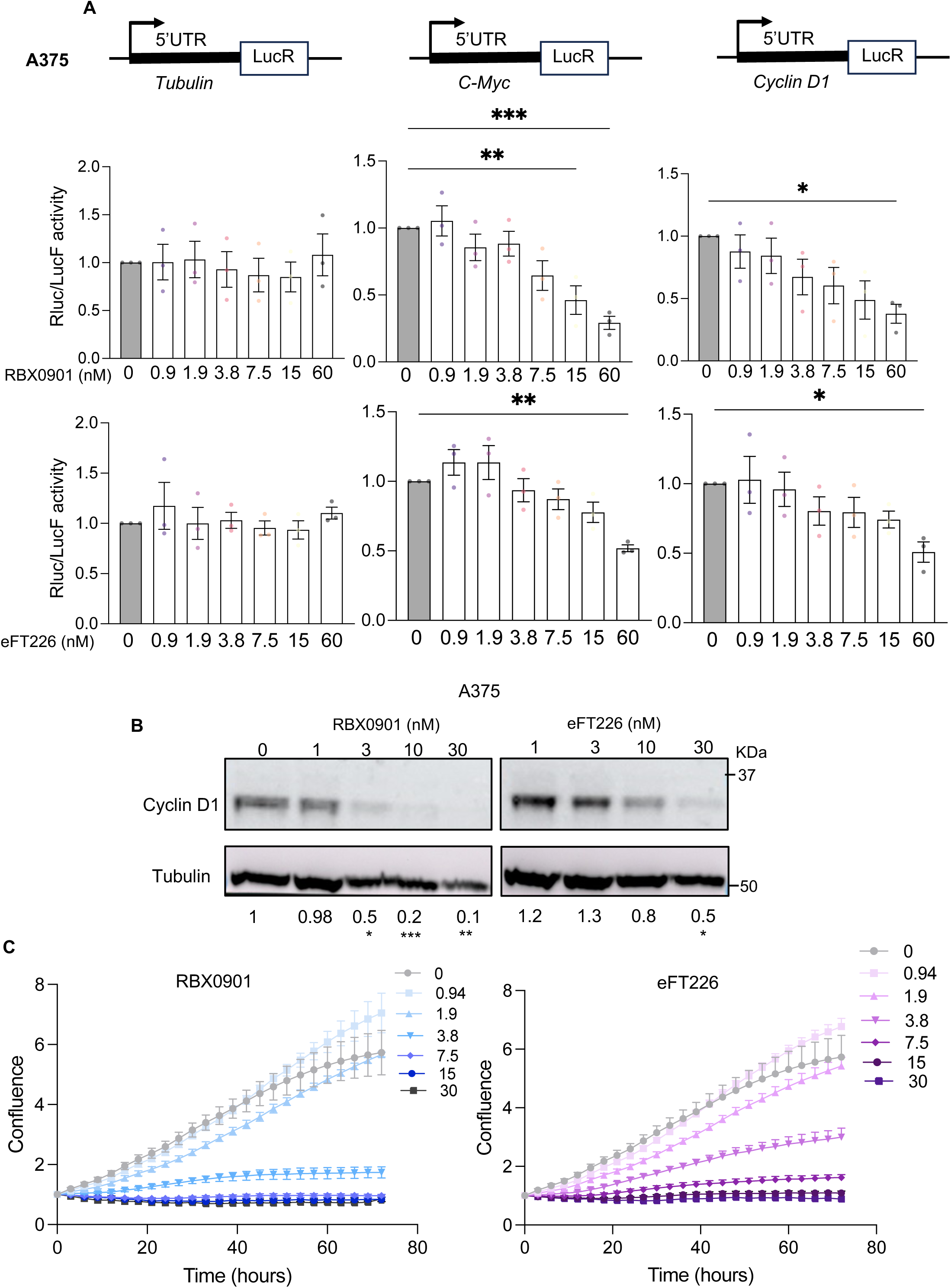
**A**, (top) Schematic representation of luciferase reporters containing the 5′UTRs of *Tubulin*, *c-MYC*, and *Cyclin D1*, used to monitor the effect of eIF4A inhibitors on reporter translation *in cellulo.* (bottom) Quantification of the luciferase assay performed in A375 cells treated with the indicated concentrations of RBX0901 or eFT226 for 24 hours. Renilla (LucR) activity was measured 48h after transfection and the activity of the Firefly luciferase (LucF) was used as control of transfection. Data are normalized to the untreated conditions and represent the mean ± SEM from 3 independent experiments (* p ≤ 0.05, ** p ≤ 0.01, *** p ≤ 0.001). **B**, Western blot illustrating Cyclin D1 protein levels in A375 cells treated with the indicated concentration of RBX0901 or eFT226 for 24 hours. Tubulin serves as a loading control, and the quantification relative to untreated conditions from 3 independent experiments is indicated. p-values were calculated by unpaired, two-tailed Student’s t-test (* p ≤ 0.05, ** p ≤ 0.01, *** p ≤ 0.001). **C**, Cell proliferation assay performed in A375 cells treated with the indicated concentrations of RBX0901 or eFT226 for 72 hours. Data represent the mean ± SEM from 3 independent experiments, and the quantification is shown relative to the untreated condition at the 72-hour time point (2-way ANOVA, **** p ≤ 0.0001).

**Supplementary Figure S9:**
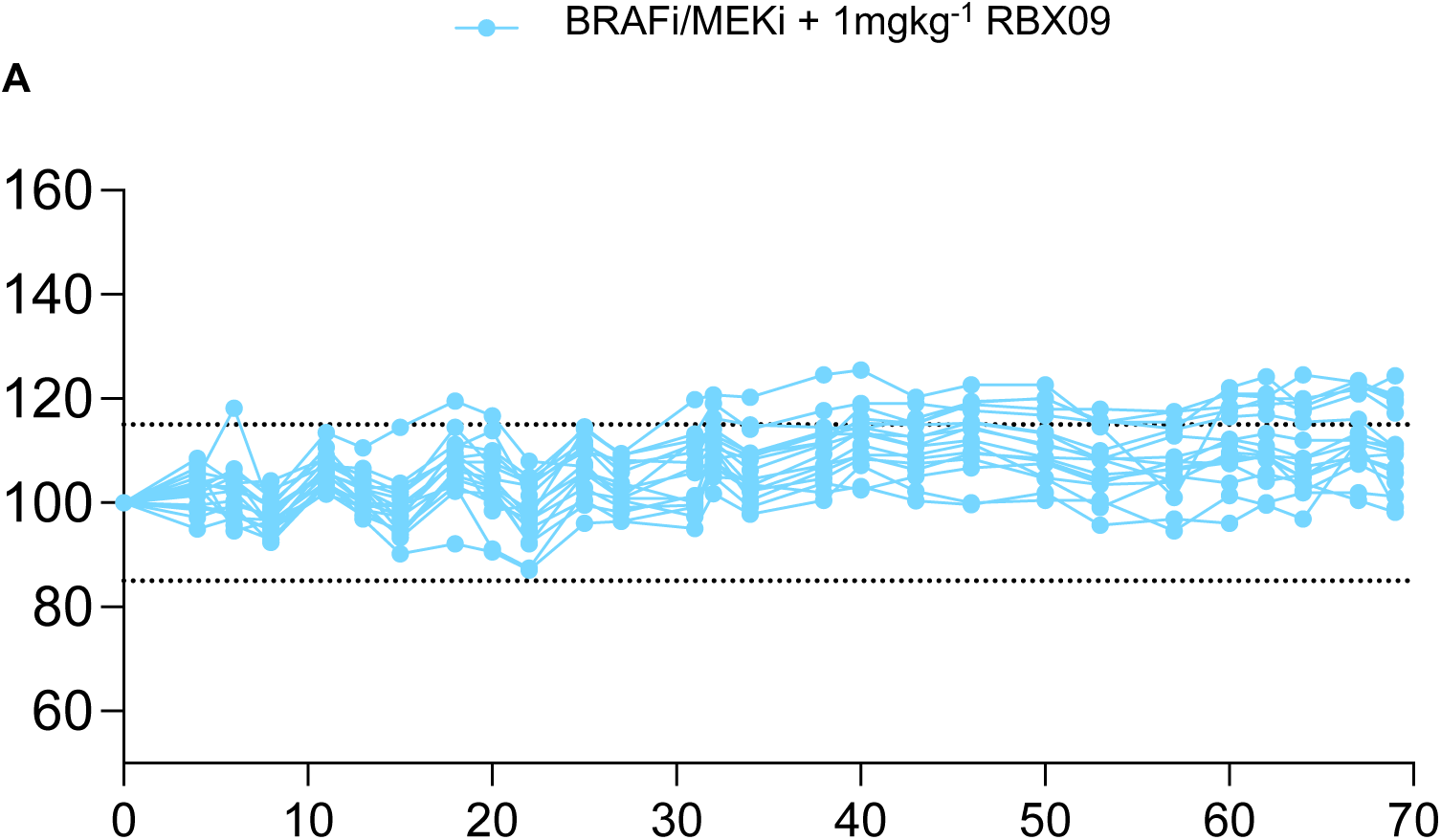
Graph showing the percentage variation in body weight of mice after 70 days of treatment with PLX4720 (BRAFi) and PD0325901 (MEKi), combined with RBX0901 (1 mg/kg).

## Notes

### Competing Interest Statement

The authors have declared no competing interest.

